# Dorsal striatal astrocytes link dopamine to slow behavioral dynamics

**DOI:** 10.1101/2024.12.01.626240

**Authors:** Iakovos Lazaridis, Emily Hueske, Gun Ahn, Wonchang Choi, Kojiro Hirokane, Sabrina Drammis, Ann M. Graybiel

## Abstract

Humans and other animals repeatedly pause from their goal-directed behavior by taking breaks. How rapid neuromodulatory signals help organize these daily engagement-disengagement cycles has remained unclear, as most studies have concentrated on in-task behavior. Here we added study of the breaks. We made simultaneous recordings of striatal dopamine release and astrocytic calcium activity as mice performed a decision-making task and took sporadic breaks. Dopamine release more strongly represented task events, whereas astrocytic calcium more strongly tracked disengagement during breaks and behavioral state cycles. Dopamine activity led to astrocytic activity, protracted adenosine signaling and dopamine depression. Blocking astrocytic calcium signals impaired engagement and rule-based learning. Astrocytes thus represent an essential aspect of the temporal organization of voluntary behavior, spanning the full engagement-disengagement cycles of daily life.

## Main Text

Our ongoing goal-directed behavior is punctuated by self-initiated breaks. These microbreaks form part of recurring cycles of engagement, disengagement, and re-engagement. Goals can be carried forward through periods of disengagement into renewed action, but how the brain accomplishes this has been unknown. Other animals also exhibit this cyclic engagement-disengagement behavior, providing an experimentally tractable and broad setting in which to study their neural organization to understand how rapid signals encoding task events during engagement can be transformed into internal trajectories that span disengagement periods and can organize the return to goal-directed behavior.

The basal ganglia are well placed to mediate this transformation (*1-4*). Striatal dopamine rapidly signals actions, outcomes, motivation, and current behavioral state (*5, 6*). Yet many changes in engagement unfold over longer timescales. Astrocytes offer a potential link between these timescales: their receptors respond to dopamine and other signaling molecules by recruiting slower calcium responses (*7-10*), and their downstream signals, such as adenosine produced by astrocytic ATP release, can prolong their effects on synapses and behavioral state (*11-14*). We asked here whether interactions between dopamine and striatal astrocytes could be central to organizing behavior across the daily cycles that lead through structured behavior interrupted by intermittent phases of disengagement behavior often unrelated to the goals followed by a return to structured action. We found that astrocytes track in remarkable detail the disengagement states so as seamlessly to bridge engagement, disengagement and re-engagement cycles in coordination with dopamine.

We examined voluntary behavior alongside dorsal striatal dopamine and astrocytic activity as mice performed a classic probabilistic choice task in a T-maze to receive rewards. They alternated spontaneously between task engagement and disengagement. Both brain signals were active across engagement states, but they carried different information: dopamine more strongly represented discrete task events during engagement, whereas astrocyte calcium more strongly represented disengagement states and progression through the engagement-disengagement cycles as a whole. Separate optogenetic stimulation of either nigral dopamine neurons or striatal astrocytes uncovered reciprocal dopamine-astrocyte interactions and also identified adenosine as a candidate downstream signal triggered by astrocytes. Suppressing astrocyte calcium as mice learned a cue discrimination learning task reduced their engagement and retarded acquisition, most strongly as the rule-driven behavior was emerging. These findings suggest that astrocyte-dopamine coupling and the extended feedback window by astrocytes are essential to organize engagement-disengagement cycles and to sustain engagement as rule-guided behaviors develop.

### Dopamine tracks trial events whereas astrocytes track engagement-disengagement dynamics

We used fiber photometry to record extracellular dopamine and astrocyte calcium simultaneously in the dorsomedial and dorsolateral striatum (DMS and DLS) across 62 sessions from five mice performing a probabilistic two-choice task with changing reward contingencies (*15, 16*) (Fig. 1, A to D and figs. S1 and S2A). Mice spontaneously paused and resumed task performance. We defined these states independent of the recorded signals by smoothing trial-initiation times into an initiation density function (IDF), which separated sustained engagement (ENG) from low-initiation disengagement (DIS) and their transitions (Fig. 1H and fig. S2, B to F). DIS included movement and isolated task actions. This allowed comparisons of task-event responses across the engagement cycle.

**Fig. 1.**
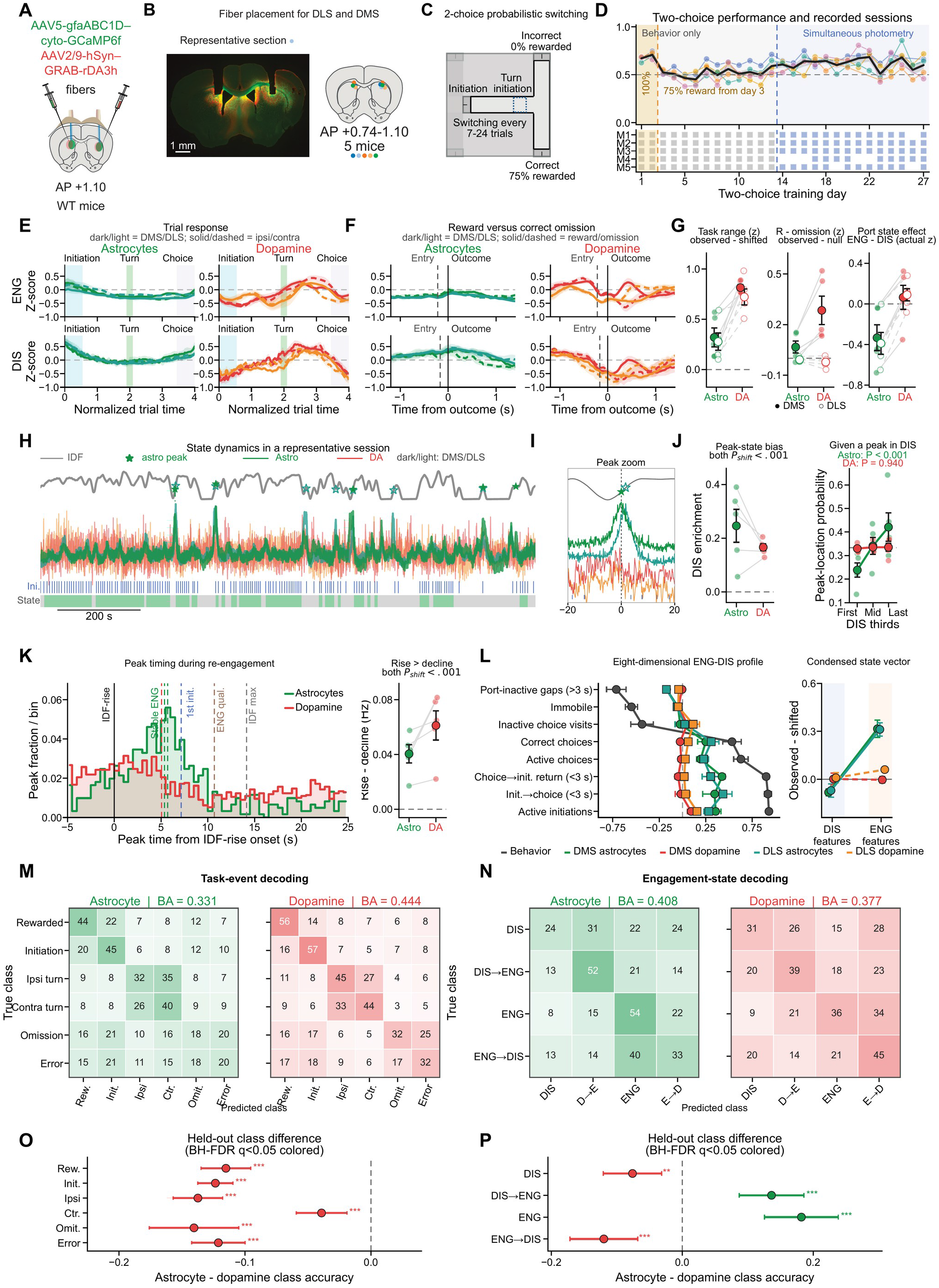
Dopamine tracks task events whereas astrocyte calcium tracks engagement dynamics. (**A** and **B**) Sensor expression and fiber placement for simultaneous extracellular dopamine and astrocyte calcium photometry in DMS and DLS. (**C** and **D**) Probabilistic two-choice task, training performance, and recording sessions (five mice; 62 sessions). (**E** to **G**) Responses to individual task events: trial-normalized profiles during ENG and DIS (**E**); outcome-aligned responses on rewarded and correct-omission trials (**F**); and task locking relative to time shifts, reward-omission contrasts relative to shuffled labels, and ENG-DIS differences in signal levels during trial-initiation and choice-port periods (**G**). (**H** and **I**) Representative session and expanded interval showing IDF, state assignments, signals, and large peaks. (**J**) Fraction of large peaks in DIS minus fraction of time in DIS (left) and peak-location probability across equal DIS thirds (right), conditional on a detected peak above the session’s 97th-percentile threshold in a complete DIS bout. Dashed line, uniform probability (one-third). (**K**) Peak timing relative to IDF rise and event-rate difference between rising and declining IDF. (**L**) Event-centered behavioral profiles compared with the ENG-DIS profile (left) and profile similarity above time-shifted controls (right). (**M** to **P**) Classification of task events and engagement states: InceptionTime held-out classification matrices (**M** and **N**) and paired astrocyte-minus-dopamine differences in held-out class accuracy with 95% confidence intervals (**O** and **P**). BA, balanced accuracy. Pale points are mice; other large symbols and error bars show mean ± SEM. Timing and label tests use within-recording randomizations; decoding comparisons use paired held-out windows. Overlapping TASK windows remain in the same cross-validation partition across signals and hemispheres. Adjusted q < 0.05, 0.01, 0.001: *, **, ***. Analysis definitions and tests: Materials and Methods, M8-M10, including M10.1. Abbreviations: Astro, astrocyte; DA, dopamine; DMS, dorsomedial striatum; DLS, dorsolateral striatum; ENG, engagement; DIS, disengagement; IDF, initiation density function; AAV, adeno-associated virus; AP, anteroposterior; WT, wild type; SEM, standard error of the mean; BH-FDR, Benjamini-Hochberg false discovery rate. Ipsi, ipsilateral; Contra/Ctr., contralateral; Ini./Init., initiation; Rew./R, reward; Omit., omission; ENG qual., engagement qualification; max, maximum. D→E and E→D denote DIS-to-ENG and ENG-to-DIS transitions; M1-M5 identify individual mice.

Within trials, dopamine activity clearly showed stronger modulation across task events than astrocyte calcium in both striatal regions, with highest dopamine activity around the turn (Fig. 1, E to G). DMS dopamine responses also differed between rewarded trials and trials in which reward was omitted after a correct choice (fig. S3, A to E). We compared signals during trial-initiation and choice-port periods between engagement states. Astrocyte calcium signals were higher during DIS than ENG, whereas dopamine signals showed little corresponding difference (Fig. 1G). This ENG-DIS difference suggested that astrocyte activity might reflect the broader range of behavioral state, beyond individual task events.

Large events were evident in the raw signals (Fig. 1H), with isolated dopamine surges preceding large astrocyte surges. We therefore asked whether the state dependence also held for the timing of these large signal events as animals progressed through disengagement and returned to engagement. We detected these events as peaks above each session’s 97th-percentile threshold. After accounting for time spent in each state, both signals were enriched during DIS. However, among peaks detected in complete DIS bouts, astrocyte peaks became increasingly likely across the first, middle, and final thirds, whereas dopamine peaks were distributed evenly (Fig. 1, H to J). Astrocyte peak timing therefore indicated a progression from entry into disengagement toward re-engagement that was not evident in dopamine peak timing. Events in both signals were more frequent around re-engagement than disengagement, so that signal changes preceded renewed task initiation (Fig. 1K and fig. S3, F to L).

To place these large signal events in a broader behavioral context, we first characterized differences in immobility, port inactivity, trial initiations, choices, and port transitions between IDF-defined ENG and DIS periods. We then compared these behavioral patterns to changes in the same measures before versus after large dopamine or astrocyte calcium events. Changes surrounding astrocyte events more closely resembled the ENG-DIS behavioral pattern than did those surrounding dopamine events in both striatal regions (Fig. 1L). We found no single task action or movement motif that accounted for this (fig. S3, M and N). Thus, large astrocyte events marked a broader behavioral trajectory through disengagement and return to the task than did dopamine.

We extended this comparison to short signal segments using InceptionTime classifiers trained to identify task events and engagement states (Materials and Methods, M10). Classifiers were tested on signal segments that had not been used for training (fig. S3, O to Q). Dopamine better identified task events, whereas astrocyte calcium better identified engagement states, particularly return from DIS and stable ENG (Fig. 1, M to P). MiniROCKET classifiers reproduced this distinction on the same test segments (fig. S3R). Astrocyte calcium therefore carried more information about the behavioral context in which performance was interrupted and resumed, while dopamine more strongly represented individual in-task events.

### Astrocytes track progression through the engagement cycle

These decoding results suggested that dopamine and astrocyte calcium carried complementary information about behavior: dopamine better identified task events, whereas astrocyte calcium better identified the breaks and state, particularly return from DIS and stable ENG. We examined whether dopamine and astrocyte calcium represented progression through successive sub-phases within ENG and DIS. For each recording, region and period around a behaviorally defined episode, we summarized event structure and ongoing activity (table S1). We generated state-phase maps with Uniform Manifold Approximation and Projection (UMAP). We could identify distinct representations of stable ENG and DIS for both astrocyte calcium and dopamine (Fig. 2A and fig. S5A). Classifiers using either signal’s features accurately distinguished these states in withheld sessions (fig. S5E). Strikingly, however, the signals differed in their organization within each state: dopamine core phases clustered closely together near the centers in both ENG and DIS states, and jumped between these, whereas astrocyte phases traced in detail successive trajectories across DIS as well as ENG (Fig. 2A). Classifiers tested on data from mice whose recordings had not been used for training also distinguished early from core phases more accurately from astrocyte than from dopamine features, particularly during ENG (fig. S5F). This organization suggested that astrocytes not only better represented disengagement than did dopamine, but more fully represented trajectories through entire ENG-DIS cycles.

**Fig. 2.**
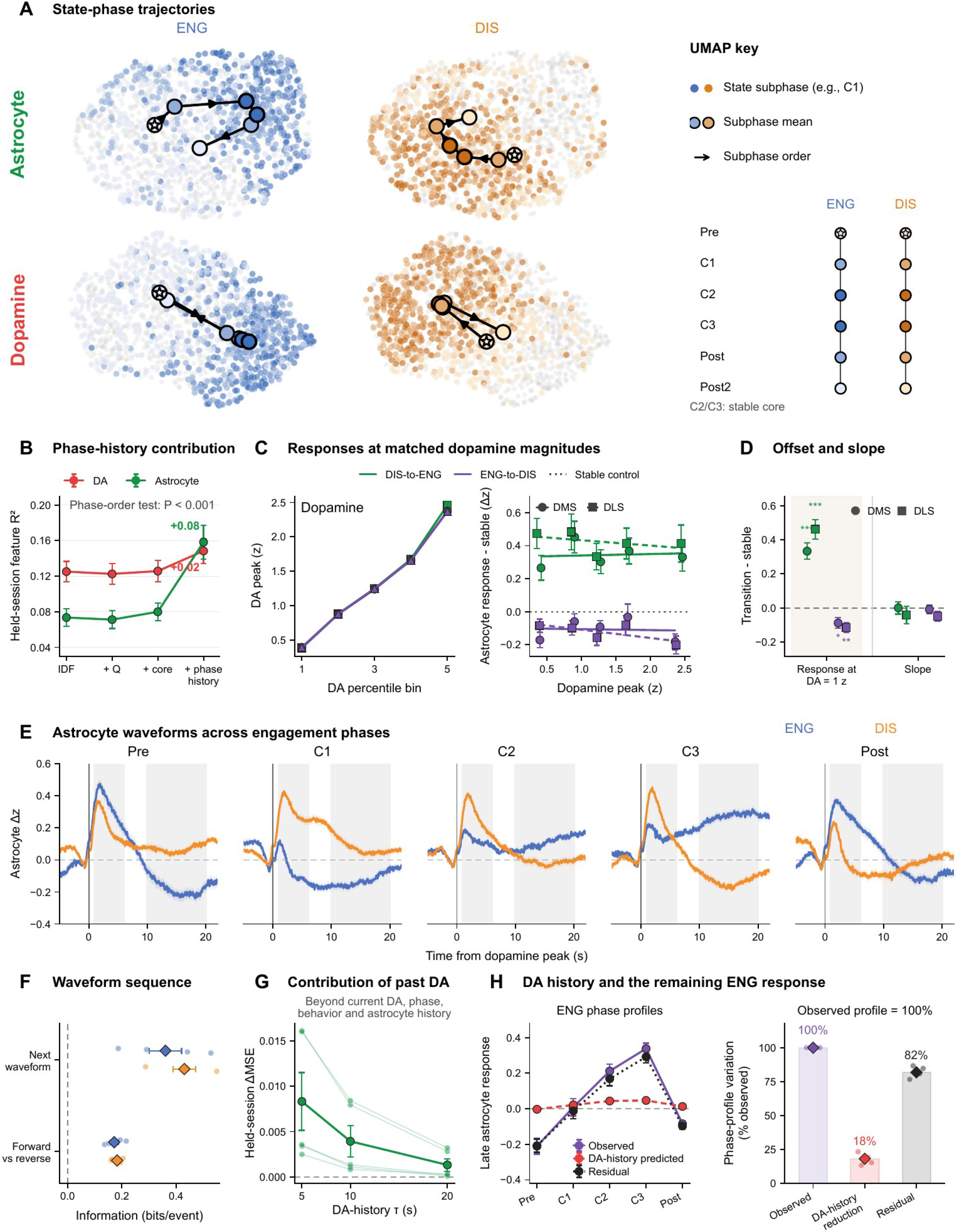
Astrocyte calcium tracks temporal progression beyond current state and recent dopamine. Five mice contributed 62 sessions with simultaneous dopamine and astrocyte calcium recordings in DMS/DLS. (**A**) UMAP trajectories of astrocyte and dopamine features during ENG and DIS. Small points are recording-session-region-phase summaries; large symbols mark phase centroids and arrows indicate temporal order. For A and B, C1-C3 are the first three quarters of a stable bout, and Post is its fourth quarter. Pre and Post2 are exterior flanks of the same duration. (**B**) Held-session feature variance explained by successive addition of IDF, fitted action value (Q), stable-core identity, and phase history. (**C**) Matched dopamine inputs and astrocyte responses. Left, mean dopamine magnitude across within-session percentile bins after equally averaging paired DMS/DLS values. Right, astrocyte transition responses minus the stable-control response within each session, region, and dopamine bin; dotted zero line, stable control. Circles/solid fits, DMS; squares/dashed fits, DLS. Green, DIS-to-ENG; purple, ENG-to-DIS. Both plots use 291 complete session-bin combinations from 61 sessions in five mice; points and error bars show session means ± SEM. Lines in the right plot are descriptive fits to bin means. Absolute regional curves are retained in fig. S5O. (**D**) Transition-minus-stable response offset at dopamine = 1 z and change in response slope, retaining the original paired regression analysis. (**E**) Dopamine-aligned astrocyte waveforms across Pre, C1-C3, and Post for ENG and DIS. Here, C1-C3 divide each stable episode into equal thirds, with equal-duration exterior Pre and Post flanks. (**F**) Predictive information in the current waveform about the next waveform beyond phase, behavior, dopamine, and region, and the advantage of forward over reversed sequence order. (**G**) Additional predictive contribution of strictly preceding dopamine history after accounting for current dopamine, phase, behavior, region, and astrocyte history. (**H**) Left, observed, dopamine-history-predicted, and residual late astrocyte responses (10-20 s after dopamine peaks) across ENG phases. Right, observed phase-profile variation normalized to 100% separately for each mouse, with mean dopamine-history reduction of 18.15% and residual of 81.85%. The quantity is the centered sum of squares across phase means, not total astrocyte variance; reduction and residual are complementary. Small points in F to H are mice; summary error bars and waveform shading show SEM. Feature definitions and contributions: table S1. Models and tests: Materials and Methods, M11 and M11.2. Additional state and response controls: figs. S5 and S6. Abbreviations: DA, dopamine; DMS, dorsomedial striatum; DLS, dorsolateral striatum; ENG, engagement; DIS, disengagement; IDF, initiation density function; UMAP, uniform manifold approximation and projection; MSE, mean squared error; SEM, standard error of the mean. ΔMSE denotes the reduction in prediction error from adding dopamine history; τ is its decay time constant, and RZ is the coefficient of determination.

We next used current IDF, action value, and ENG or DIS identity to identify the signal representations, and then added temporal position within the ordered phases of each state (phase history) to quantify its additional contribution. Current IDF made the largest predictive contribution for dopamine, whereas phase history contributed most for astrocyte calcium (Fig. 2B). If so, astrocyte responses to dopamine might therefore depend on position within the engagement cycle. We tested this by examining how the relationship between the two signals varied with state and phase, using measurements of relative timing, covariation, and event-aligned responses. Together, these measurements distinguished ENG from DIS and early from core phases within ENG (fig. S5, A to F and H to I). Thus, temporal progression was reflected in astrocyte activity and its relationship to dopamine, whereas dopamine activity was more closely associated with current initiation density.

Astrocyte responses also differed across phases even when preceding dopamine magnitudes were matched: dopamine-aligned astrocyte responses were enhanced during return to engagement and reduced during departure from engagement relative to stable-state responses in both DMS and DLS (Fig. 2C and fig. S5O). The difference appeared primarily as an offset in response magnitude, without a detectable change in its scaling with dopamine (Fig. 2D). Relative astrocyte recruitment was greatest before ENG entry, and response waveforms evolved through successive phases of both ENG and DIS (Fig. 2E and figs. S5, K to O, and S6, A to C and F to I). But phase-dependent modulation also extended to later parts of the dopamine-aligned response, as shown by measurements of astrocyte activity in early (1-6 s) and later (10-20 s) windows after the dopamine event. For these response trajectories, we divided each identified stable ENG or DIS episode into three equal-duration periods, labeled C1, C2, and C3 from beginning to end. Mean late activity rose from below the pre-event baseline in C1 to above it in C3 during ENG, with the opposite progression during DIS (Fig. 2E and fig. S6, B, F, and G). Measured task events and movement did not fully account for this organization (fig. S5, K to N). Astrocyte activity increased during the IDF-defined return to engagement before the first renewed trial initiation (Fig. 1K and fig. S3, J to L), and responses to the same initiation action varied with cycle position (fig. S6, H and I). Thus, similar dopamine events could accompany different astrocyte responses according to behavioral progression.

We next examined whether these evolving responses followed a predictable temporal order. We defined recurring dopamine-aligned astrocytic waveform patterns in training sessions and tested their predictive information in withheld sessions. The current pattern predicted the next pattern beyond exact behavioral phase, current behavior, and dopamine magnitude; prediction also favored the chronological sequence over its reverse (Fig. 2F and fig. S6J). Prediction of upcoming transitions between phases of the engagement cycle by astrocyte activity was more selective. Preceding activity was most informative at DIS entry and early progression, whereas current waveforms were informative at later ENG boundaries (fig. S6, K and L). Thus, astrocyte responses carried information about subsequent waveforms and selected, rather than all, behavioral transitions (fig. S6M).

Recent dopamine improved prediction of astrocyte responses within a known phase, beyond current dopamine, movement, and preceding astrocyte activity, in models tested on withheld sessions (Fig. 2G and fig. S6D). Preceding astrocyte activity also improved prediction of the subsequent dopamine-aligned astrocyte response beyond dopamine, immediate behavior, and state/history predictors (fig. S5J). Across phases, however, temporal position accounted for more variation in astrocyte responses than in current or recent dopamine activity (fig. S6E). To assess dopamine’s contribution across phases, we modeled the astrocyte trajectory using recent dopamine without phase labels. The model reproduced the shape of the late astrocyte response to dopamine peaks (10-20 s) across ENG phases, but removing the modeled dopamine-history component reduced variation across phase means only by about 18%, leaving about 82% of the observed phase-profile variation (Fig. 2H). Dopamine history therefore explained some within-phase variation but only a limited part of the broader astrocyte trajectory, consistent with there being additional state-related inputs.

### Dopamine and astrocytes interact across timescales

During behavior, dopamine events preceded increased astrocyte activity, and astrocyte activity preceded a later reduction in dopamine (fig. S4). This suggested reciprocal interactions, but recordings alone could not establish either causal direction. We tested each direction by stimulating dopamine neurons or striatal astrocytes in complementary experiments.

We expressed ChrimsonR opsin in dopamine-containing neurons of the substantia nigra pars compacta (SNc) in DAT-Cre mice and stimulated their cell bodies with 4-s light trains. With fiber photometry probes in the striatum, we measured striatal dopamine, astrocyte calcium, or adenosine in separate mouse cohorts in response to the stimulation. We included adenosine as a candidate downstream signal through which dopamine-evoked astrocyte recruitment could influence striatal circuit activity over longer timescales (*7, 17-20*). The dopamine and astrocyte recordings sampled both DMS and DLS (Fig. 3, A to C). Striatal dopamine peaked first, followed by astrocyte calcium and then adenosine. Adenosine decayed much more slowly than either dopamine or astrocyte calcium, remaining elevated for tens of seconds after both other responses had subsided. These experiments demonstrated that a brief nigral dopaminergic input could recruit more persistent downstream signaling in the striatum.

**Fig. 3.**
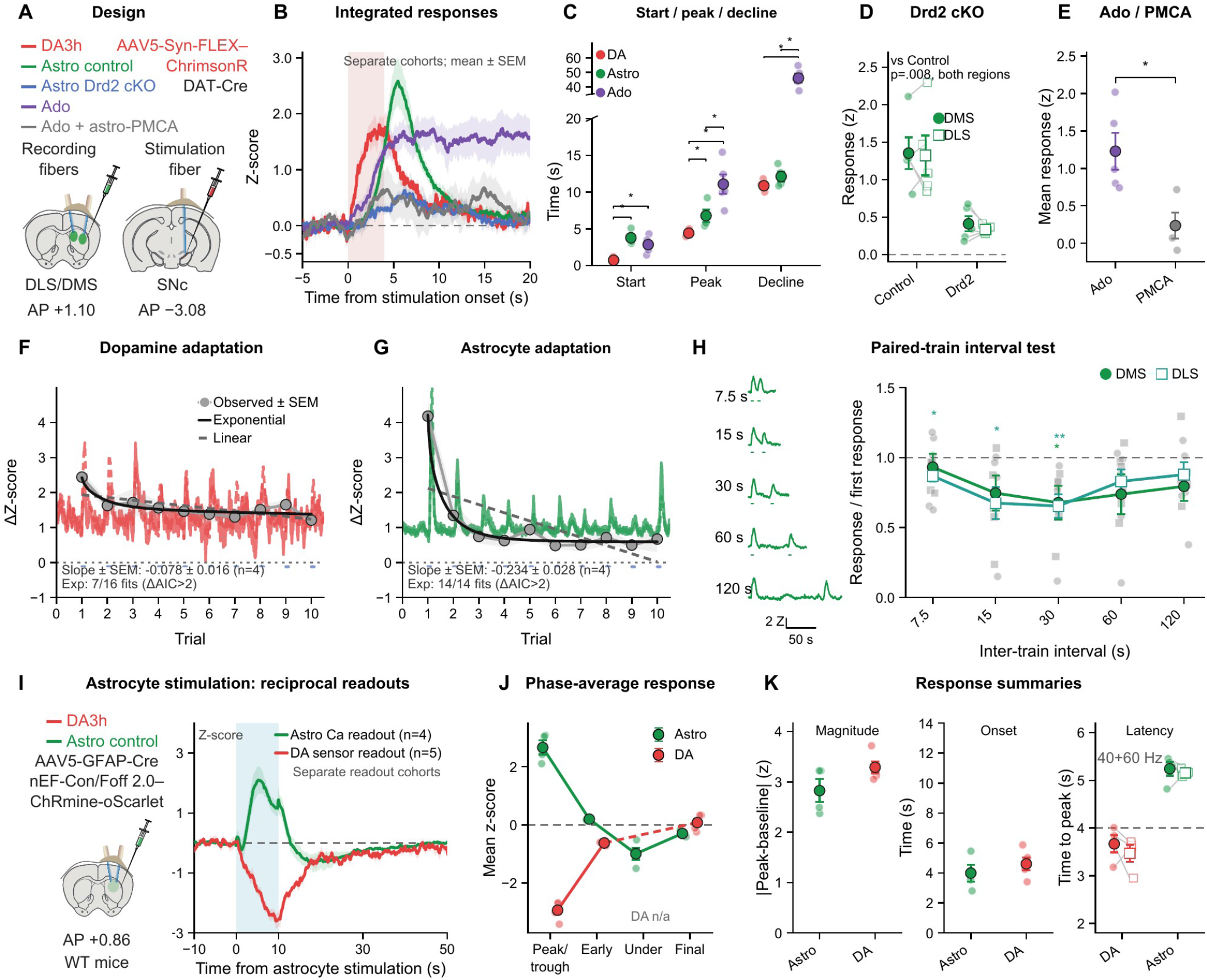
Dopamine recruits slower astrocyte and adenosine responses, and astrocyte activation suppresses dopamine. (**A** to **C**) Response timescales after dopamine-neuron stimulation. ChrimsonR stimulation of SNc dopamine-neuron cell bodies with fiber photometry of striatal dopamine, astrocyte calcium, or adenosine in separate sensor cohorts (**A**). Responses aligned to a 4-s, 40-Hz stimulation train; astrocytic *Drd2* deletion and PMCA expression are included for calcium and adenosine, respectively (**B**). Response onset, peak, and decline times (**C**); adenosine persists for tens of seconds after dopamine and calcium decline. (**D**) Evoked astrocyte calcium magnitude in control and astrocytic *Drd2*-deletion mice. (**E**) Evoked adenosine magnitude with or without astrocytic PMCA. (**F** to **H**) Dependence on recent stimulation. Responses across ten identical trains: time courses, per-train magnitudes, and attenuation fits (four mice per sensor cohort) (**F** and **G**). Astrocyte responses decline more rapidly than dopamine responses. Second-to-first astrocyte response ratios at different paired-train intervals (**H**); dashed line, equal responses. (**I** and **J**) Dopamine responses to astrocyte activation. Dorsal-striatal astrocyte stimulation with ChRmine and aligned calcium or dopamine responses (**I**). One fiber delivered 10 s of continuous light; a second recorded from the same striatal volume in separate calcium (n = 4 mice) and dopamine (n = 5) cohorts. Mean responses around the peak/trough, early recovery, calcium undershoot, and final baseline return (**J**); no undershoot phase is defined for dopamine. (**K**) Absolute response magnitude (left) and onset latency (middle) after astrocyte activation. The right plot shows dopamine and astrocyte peak latencies after SNc dopamine-neuron stimulation, averaged across 40 and 60 Hz. Circles and squares denote DMS and DLS; the dashed line marks the end of the 4-s train. Time zero denotes stimulation onset; shaded stimulus intervals mark light delivery. Small points are subject summaries; lines, large symbols, and error bars or response shading show mean ± SEM. Cohort-specific stimulation protocols, response definitions, sample sizes, and statistical tests: Materials and Methods, M1.1, M2.1-M2.2, M2.5, and M13. Abbreviations: Astro, astrocyte; DA, dopamine; Ado, adenosine; Ca, calcium; DMS, dorsomedial striatum; DLS, dorsolateral striatum; SNc, substantia nigra pars compacta; cKO, conditional knockout; Drd2, dopamine D2 receptor; PMCA, plasma-membrane calcium ATPase; DAT, dopamine transporter; GFAP, glial fibrillary acidic protein; AAV, adeno-associated virus; AP, anteroposterior; WT, wild type; SEM, standard error of the mean; AIC, Akaike information criterion; Exp, exponential fit; Under, undershoot; n/a, not applicable. DA3h denotes the dopamine sensor.

We made astrocyte-specific receptor deletions of dopaminergic and noradrenergic receptors and identified a strong D2-receptor contribution to this recruitment. Under this protocol, *Drd2* deletion reduced evoked astrocyte calcium by approximately 70-75% in DMS and DLS, whereas *Drd1* or *Adra1a* deletion produced no detectable reduction (Fig. 3D and fig. S7, C and D). We then tested the dependence of the prolonged adenosine response on astrocyte calcium by expressing plasma-membrane calcium ATPase (PMCA) selectively in striatal astrocytes to enhance calcium extrusion and deplete it from the astrocytes (*21*). PMCA reduced evoked adenosine by approximately 80% relative to controls (Fig. 3E and fig. S7E). These interventions identify astrocytic D2 receptors as a major route for dopaminergic recruitment and astrocyte calcium as a major mediator of prolonged extracellular adenosine.

Recent dopaminergic input also altered subsequent astrocyte recruitment under controlled stimulation, consistent with the history dependence observed during behavior. Across ten identical SNc-stimulation trains, with repeated responses averaged within mice, astrocyte responses attenuated approximately three times faster than dopamine responses, particularly over the first few trains (Fig. 3, F and G, and fig. S7F). This change could not be explained simply by weaker dopamine input. We assessed the timescale of this dependence by varying the interval between trains in a separate paired-train experiment. The second-to-first astrocyte response varied with the interval in DMS and DLS over tens of seconds (Fig. 3H).

For the reciprocal test, we expressed ChRmine in dorsal-striatal astrocytes and delivered 10 s of continuous light while recording astrocyte calcium or dopamine from the same striatal volume, in separate cohorts of, respectively, four and five mice. The astrocyte stimulation evoked a calcium elevation and delayed, prolonged, reversible dopamine suppression (Fig. 3, I to K). Synchronized open-field tracking detected no consistent stimulation-locked movement changes over these time periods (fig. S7, J to M). Astrocyte activation thus regulates subsequent striatal dopamine over an interval extending beyond the initial calcium response.

These perturbations establish reciprocal interactions: dopamine history alters astrocyte recruitment, and astrocyte activation suppresses subsequent dopamine. The prolonged, calcium-dependent adenosine response provides a candidate mechanism for extending the interaction across time. Mechanisms underlying this dopamine suppression remain to be identified.

### Astrocyte calcium supports engagement during learning

Given the close tracking of the engagement cycles by the striatal astrocytes and their reciprocal interactions with dopamine, we hypothesized that astrocyte calcium could contribute to engagement during reinforcement learning. We expressed PMCA bilaterally in striatal astrocytes in seven mice to block astrocytic calcium and compared their behavior with that of eight GFAP-Cre-injected controls. The viral expression obtained covered DMS and DLS, most strongly DMS (Fig. 4, A and C and fig. S8, A and B). In the head-fixed task we used, auditory cues predicted sucrose reward or aversive light. Licking during a designated pre-delivery response period increased sucrose amount in reward trials and light intensity in cost trials (Fig. 4B). We quantified cue discrimination with d′ and response bias with C, a signal-detection measure of the tendency to respond across cue types (*22*). We use −C as an engagement measure here, with higher values indicating greater responding, distinct from initiation-based engagement in the recording task. Both groups developed reward-cost selectivity (Fig. 4D), enabling engagement comparisons across discrimination stages.

**Fig. 4.**
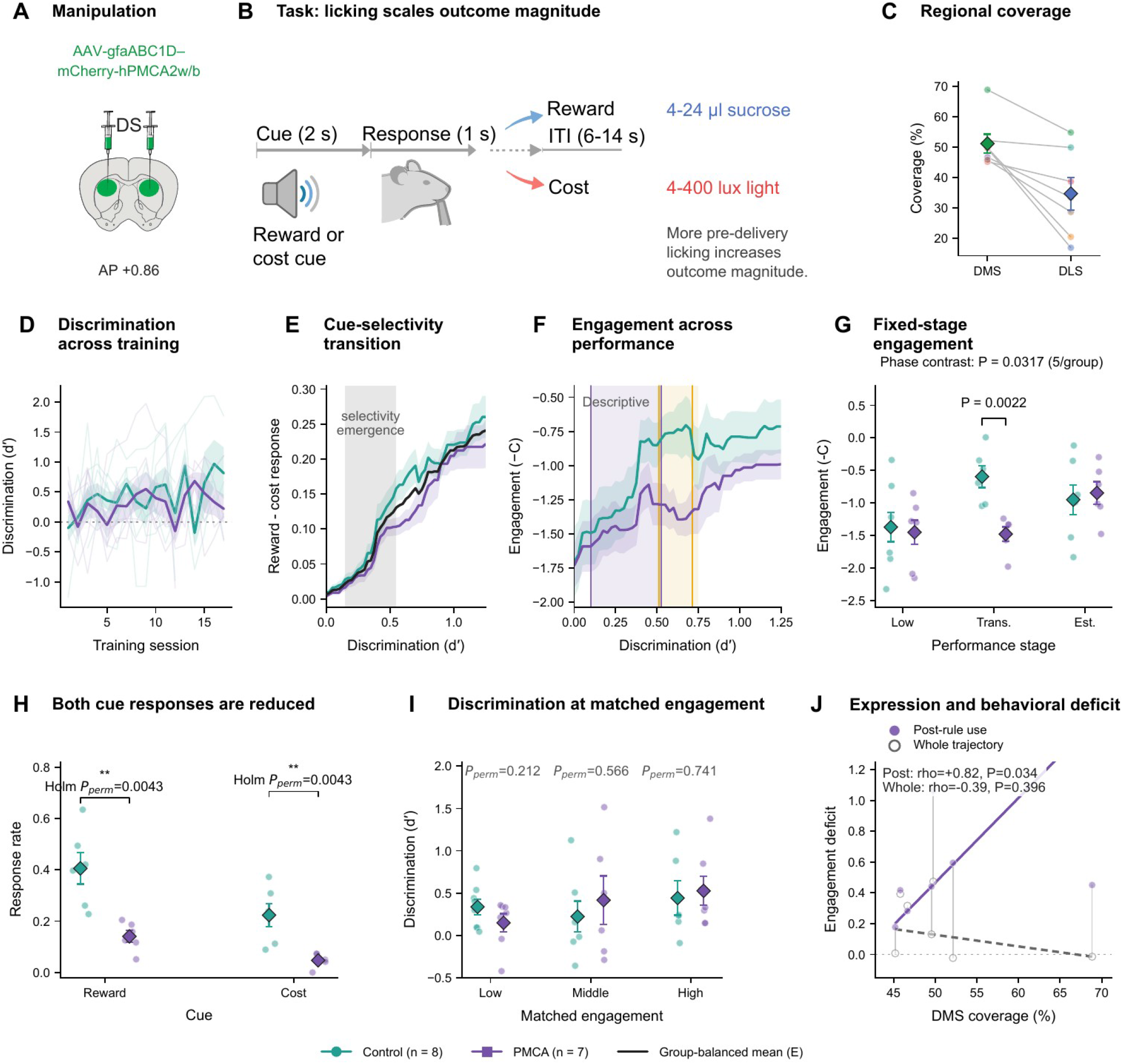
Suppressing astrocyte calcium reduces engagement as cue-guided behavior emerges. (**A**) Bilateral dorsal-striatal expression of astrocytic hPMCA2w/b. (**B**) Auditory approach-avoidance task. Licking in the designated pre-delivery response period increased sucrose reward (4-24 µl) or aversive-light magnitude (4-400 lux); withholding that response did not prevent outcome delivery. (**C**) Paired DMS/DLS expression coverage within mice. (**D**) Discrimination (d′) across training sessions. (**E**) Reward-minus-cost response probability across discrimination levels, with the group-balanced curve locating the emergence of cue-selective responding. (**F**) Engagement (−C) across discrimination levels, showing lower responding by PMCA mice around intermediate performance. In D-F, teal denotes GFAP-Cre controls and purple denotes PMCA mice; the black curve in E is the group-balanced mean. (**G**) Engagement during fixed low, transition (d′ = 0.5-0.75), and established performance stages. (**H**) Reward- and cost-cue response probabilities during transition; PMCA mice responded less to both cues. The joint response geometry is retained in fig. S8G. (**I**) Discrimination at matched low, middle, and high engagement. (**J**) Association between DMS expression coverage and engagement deficit, calculated after cue selectivity emerged or across the whole trajectory. Greater coverage accompanied a larger deficit after selectivity emerged. Teal, GFAP-Cre controls (n = 8 mice); purple, PMCA (n = 7); six mice per group contributed to transition comparisons, and five per group with measurements at all three stages contributed to the phase contrast in G. Pale points or traces are mice; large symbols, error bars, and shading show mean ± SEM. The overlapping-window trajectory in F is descriptive; inferential comparisons use the fixed stages. Regional expression and behavioral controls are in figs. S8 and S9. Statistical tests and behavioral and expression analyses: Materials and Methods, M3.1 and M14. Abbreviations: DS, dorsal striatum; DMS, dorsomedial striatum; DLS, dorsolateral striatum; AP, anteroposterior; ITI, inter-trial interval; AAV, adeno-associated virus; GFAP, glial fibrillary acidic protein; PMCA, plasma-membrane calcium ATPase; hPMCA2w/b, human PMCA2 w/b variant; SEM, standard error of the mean; Trans., transition; Est., established. C is the signal-detection decision criterion; Pperm denotes a permutation-test P value, and rho denotes Spearman’s rank correlation coefficient.

Engagement was most strongly reduced in PMCA mice during the transition between low and established discrimination, when they lagged in differential responding and instead responded less to both reward and cost cues (Fig. 4, E to H and fig. S8G). This deficit exceeded the average group difference in the low-discrimination stage before the transition and the established-discrimination stage after it (Fig. 4G and fig. S8, E to H). Reduced responding to both cues indicates a change in overall response propensity, rather than simply a shift in reward-cost preference. The stage dependence of this engagement deficit indicated that astrocyte calcium was especially important as animals began to implement the emerging cue rule.

At matched levels of engagement (−C), no difference in cue discrimination was detected between PMCA and control mice (Fig. 4I and fig. S8, I to K). Response licking and consummatory licking were maintained, and the same mice had shown no broad impairment on simpler operant tasks (figs. S8, L and M, and S9). These controls argue against a general inability to lick, consume reward, or perform rewarded actions. Fewer PMCA mice nevertheless reached high discrimination (fig. S8N), which could reflect reduced engagement, an additional learning effect, or both. The clearest behavioral effect was reduced engagement as cue-guided responding developed.

The anatomy bolstered this result: the greater the DMS expression coverage in individual PMCA mice, the larger the engagement deficit after cue selectivity emerged (Fig. 4J). The viral expression also involved DLS; this association supports a DMS-weighted contribution, but leaves open additional effects from DLS. Collectively, these experiments support a contribution of dorsal-striatal astrocyte calcium to engagement during cue-guided learning.

## Discussion

Our findings identify an ordered representation of engagement-disengagement cycles in the activity patterns of astrocytes in the dorsal striatum. They, rather than striatal dopamine, predominate in tracking breaks: the astrocyte calcium tracked entry into, progression through, and exit from disengagement, distinguishing where the animal was within the behavioral cycle. In direct comparisons of striatal dopamine and astrocytic calcium activity, dopamine more strongly represented discrete task events, whereas astrocyte calcium more strongly represented engagement state and its successive phases. This trajectory information appeared in both continuously recorded astrocyte signals and in dopamine-aligned astrocyte responses. We found no single measured behavior, task event, or movement pattern that could account for this temporal organization, supporting modulation by progression through the engagement-disengagement cycle. The astrocyte activity that we identify here therefore adds an ordered temporal dimension to the striatal representations of behavior, extending from individual events to the organization of breaks and renewed engagement.

The relationship between astrocytic activity and dopamine depended on engagement state, temporal phase, and preceding dopamine activity. Astrocyte responses scaled with local dopamine magnitude, with recent exposure accounting for additional within-phase variation. Yet comparable dopamine events elicited different responses across the engagement cycle, particularly in the delayed component. Further, dopamine features and recent dopamine history together recovered less of the ordered state trajectory than did astrocyte features. These observations suggest that astrocytes incorporate dopaminergic input within a broader representation of behavioral context and offer full coverage of cyclical bouts of engagement and disengagement. The identification of temporal trajectories tracked through cyclical engagement-disengagement cycles by astrocytes is the key component discovered here, with a modest but important contribution by dopamine. Local network activity and other state-related inputs may make responses to individual dopamine events depend on position within the engagement cycle.

Our findings support a working sequence linking rapid striatal dopamine events to slower reciprocal signaling with astrocyte activity. Dopamine recruits astrocyte calcium through astrocytic *Drd2*; recent dopamine exposure and circuit state shape the response to each new event; and astrocyte purinergic output, identified here by adenosine recordings, opens a slower feedback window that can potentially restrain subsequent dopamine release. Neuromodulator-dependent gating in Drosophila and cultured mammalian astrocytes provides a precedent for context-dependent recruitment (*23*). Our prolonged adenosine response and astrocyte-evoked dopamine trough suggest feedback analogous to A1-mediated inhibition of dopamine release in the nucleus accumbens (*17*). Adenosine signaling and ENT1 also link dorsomedial-striatal astrocyte activation to a shift from habitual to goal-directed behavior in mice (*18*). We note that preferential A2A receptor expression by indirect-pathway D2 spiny projection neurons (SPNs) suggests that the astrocyte-dependent adenosine could also bias the balance between direct- and indirect-pathway activity (*24, 25*). Such biasing could help reorganize circuit state and task drive during disengagement, before renewed engagement and action.

Compartmental differences in adenosine production could provide a further level of regulation. High enrichment of NT5e fluorescence in the matrix compartment in the striatum relative to MOR1-rich striosomal zones (fig. S7, G to I), together with earlier evidence for compartmental 5′-nucleotidase activity (*26*), suggests a spatial basis for this possibility. In light of striosomal pathways controlling nigral dopamine neurons (*16*), astrocyte-dependent adenosine signaling could help regulate to striosome-matrix interactions and their influence on dopamine.

We show here that astrocyte calcium also contributed to engagement as acquisition of reinforcement cue-guided behavior developed. Suppressing astrocyte calcium most strongly reduced cue-response propensity during the critical learning transition when an emerging rule began to guide behavior. This finding connects astrocyte activity to behavioral participation during the development of acquired task structure. This timing suggests that astrocyte function may be particularly relevant when recent experience must organize subsequent action. The mechanism linking the recorded state trajectories to this behavioral effect remains to be established. Together, however, our recording and perturbation results motivate a model in which the temporal organization of astrocyte activity contributes to maintaining engagement while behavior is being shaped by learning.

Human studies provide examples of potential functions for breaks between periods of work or practice. A meta-analysis of microbreaks found small benefits for vigor and fatigue but no significant overall performance benefit (*27*). Brief, rare task-goal switches can prevent vigilance decline (*28*), and short rests between motor-practice blocks are associated with skill gains (*29*) and hippocampo-neocortical replay that predicts those gains (*30*). Whether spontaneous disengagement in the present task supports recovery or memory processing remains to be tested.

More broadly, our findings place microbreaks within the temporal organization of voluntary behavior. Studies of striatal function have largely emphasized actions, outcomes, and learning during task performance (*5, 6, 31-33*). Extending this view across breaks in engagement brings the continuity between successive periods of engagement into focus. Astrocytes, whose calcium dynamics and downstream signaling extend beyond individual neuronal events, are suited to this temporal role. They become the foundation of cyclical encoding of disengagement as well as the entire cycles of engagement-disengagement states across time. It is known that astrocytes contribute to circadian timekeeping in the suprachiasmatic nucleus (*34*), and that hypothalamic astrocytic insulin signaling links metabolic cues to circadian behavior through sex-dependent dopamine signaling in mice (*35*). Striatal astrocyte manipulations can alter behavioral flexibility and peripheral metabolism in male mice (*14, 36*), and astrocytic adenosine A2B signaling supports brain energy metabolism, synaptic function, and sleep (*37*). Striatal astrocytes are also related to habits and stereotypies (*18, 21*), which place relatively low demands on in-task encoding (*38*) but maintain organization across engagement-disengagement cycles. An intriguing possibility is that the irregularly occurring but highly cyclical behavioral cycles of engagement-disengagement that we here show to be tracked by astrocytes relate mechanistically to the highly regular, longer rhythms of astrocyte function related to circadian, metabolic, and sleep-wake regulation. By tracking progression through engagement and disengagement states and integrating recent dopamine activity, dorsal striatal astrocytes could link rapid neuromodulatory events to the extended organization of voluntary behavior across disengagement and the return to engagement and goal-directed action.

## Supplementary Results

### 1 Task events and the engagement cycle

#### Recording sites, behavioral organization, and signal robustness

**Fig. S1.**
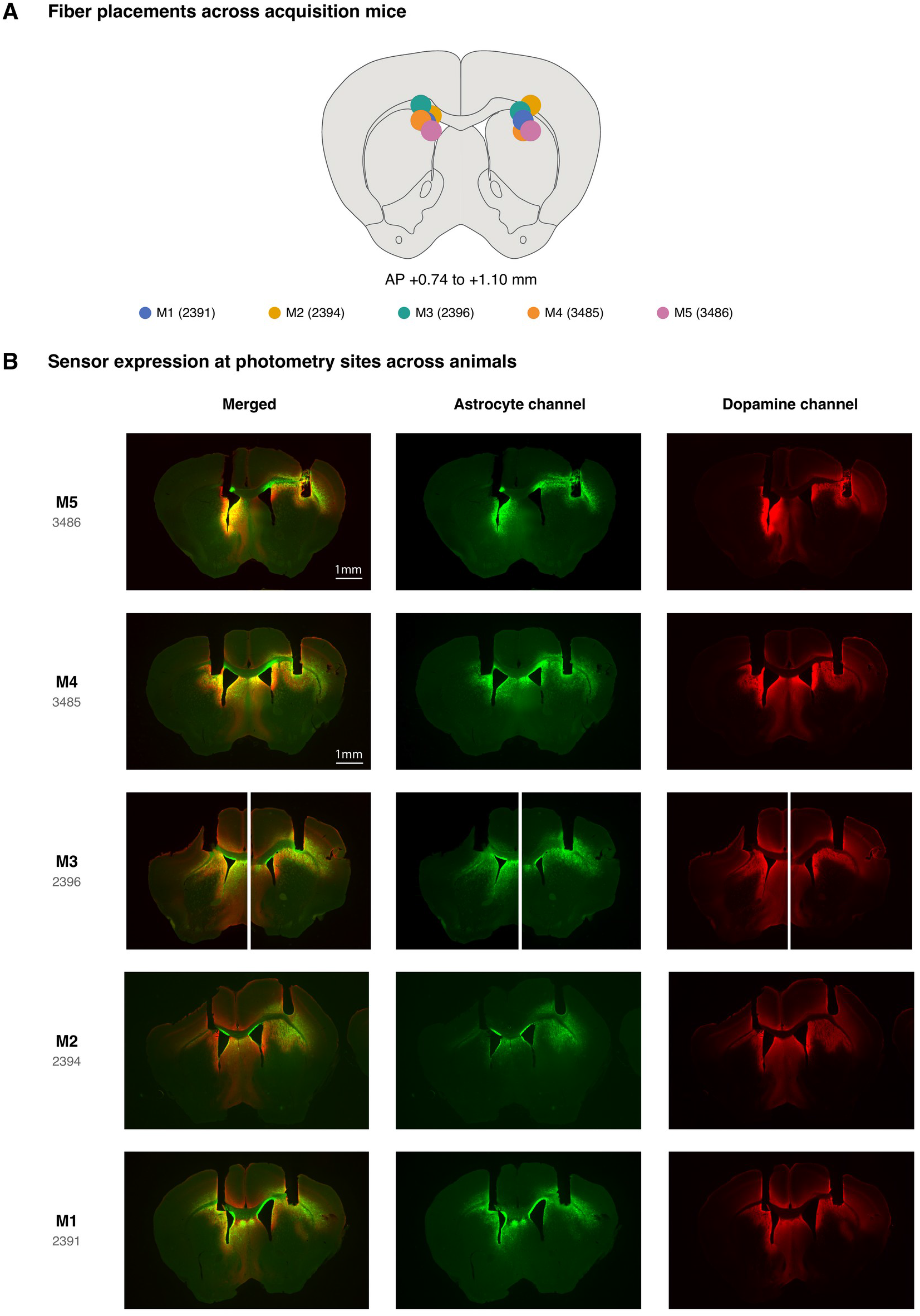
Sensor expression and fiber placement in the five recording mice. (A) Fiber-tip locations in the right dorsolateral striatum (DLS) and left dorsomedial striatum (DMS). (B) Matched coronal sections show astrocyte-calcium and dopamine-sensor expression beneath the fiber tips in each mouse. Merged and individual channels verify sensor expression at the photometry sites sampled in Figure 1. Scale bars, 1 mm. Sensor-expression and fiber-placement procedures: Methods, M2.3 and M3.

Histology confirmed dopamine- and astrocyte-sensor expression at the photometry sites in the right DLS and left DMS of all five mice (fig. S1; Methods, M2.3 and M3).

The fitted action-value estimate tracked rewarded-side changes and observed choices (fig. S2A). Independently, trial-initiation times identified sustained ENG and DIS periods without using either recorded signal (fig. S2, B and C). Mice retained the task’s movement geometry during DIS but took longer from initiation to choice (fig. S2D). Within rewarded-side blocks, accuracy and estimated action value increased while IDF and ENG probability decreased (fig. S2, E and F). Engagement therefore described the organization and frequency of task participation, which could decline even as choices became more accurate. State definitions and block analyses are described in Methods, M8 and M10.2.

Task-response differences were preserved across preprocessing choices. The selected 180-s baseline removed 76.8% of aggregate absolute drift while retaining the overall trace structure (Pearson r=0.880 ± 0.008; fig. S2, G and H). Constructed events retained their peaks without measurable artificial undershoot, and task-response profiles remained similar across baseline durations (fig. S2, H and I; Methods, M5.3).

#### Reward timing and port departure

Reward delivery and port departure occurred close together, raising the possibility that the delayed DMS dopamine component reflected departure timing. Separating trials by reward-to-exit interval distinguished these alternatives: the reward-minus-omission component remained at a similar latency after reward but shifted relative to port exit (fig. S3, A and B). Reward alignment was tighter in four of five mice, and peak latency relative to reward changed little across interval groups (fig. S3, C and D). The DMS dopamine contrast also exceeded the reward-label shuffle (fig. S3E; Methods, M10.3), supporting an outcome-related component rather than a consequence of variable port-exit timing.

#### Robustness and behavioral context of large signal events

DIS enrichment persisted across the 95th-, 97th-, and 99th-percentile peak thresholds, after accounting for unequal state duration, and under alternative behavior-only state definitions (fig. S3, F to H). Dividing bouts into quarters also resolved the changing astrocyte event rate through DIS and its decrease during ENG in both regions (fig. S3I).

Within-bout permutation tests supported the different timing profiles described in the main Results: astrocyte peaks were nonuniformly distributed through DIS (P<0.001), whereas dopamine peaks were not (P=0.940; Methods, M10.1).

Transition-related signal changes were also present in continuous recordings without selecting large events. Relative to matched stable-DIS periods, signals increased around the onset of the behavior-defined IDF rise (fig. S3, J and K). Signal maxima preceded the first renewed initiation in all five mice for dopamine and four of five for astrocyte calcium (fig. S3L; Methods, M10.6).

Dopamine peaks were enriched around turns, whereas astrocyte peaks showed smaller, distributed associations with task actions (fig. S3M). Astrocyte-event enrichment also varied across movement labels and animals; no single measured task action or movement motif accounted for the astrocyte-event pattern (fig. S3N).

#### Decoder validation

The InceptionTime task comparison was consistent across held-out folds despite unequal class counts; state-decoding performance varied more across folds (fig. S3, O to Q). MiniROCKET reproduced the complementary information carried by the signals using the same test observations: dopamine gave higher task accuracy (0.373 versus 0.300), whereas astrocytes gave higher state accuracy (0.416 versus 0.367; fig. S3R). Thus, the distinction was reproduced across classifier architectures (44, 46; Methods, M10.4).

**Fig. S2.**
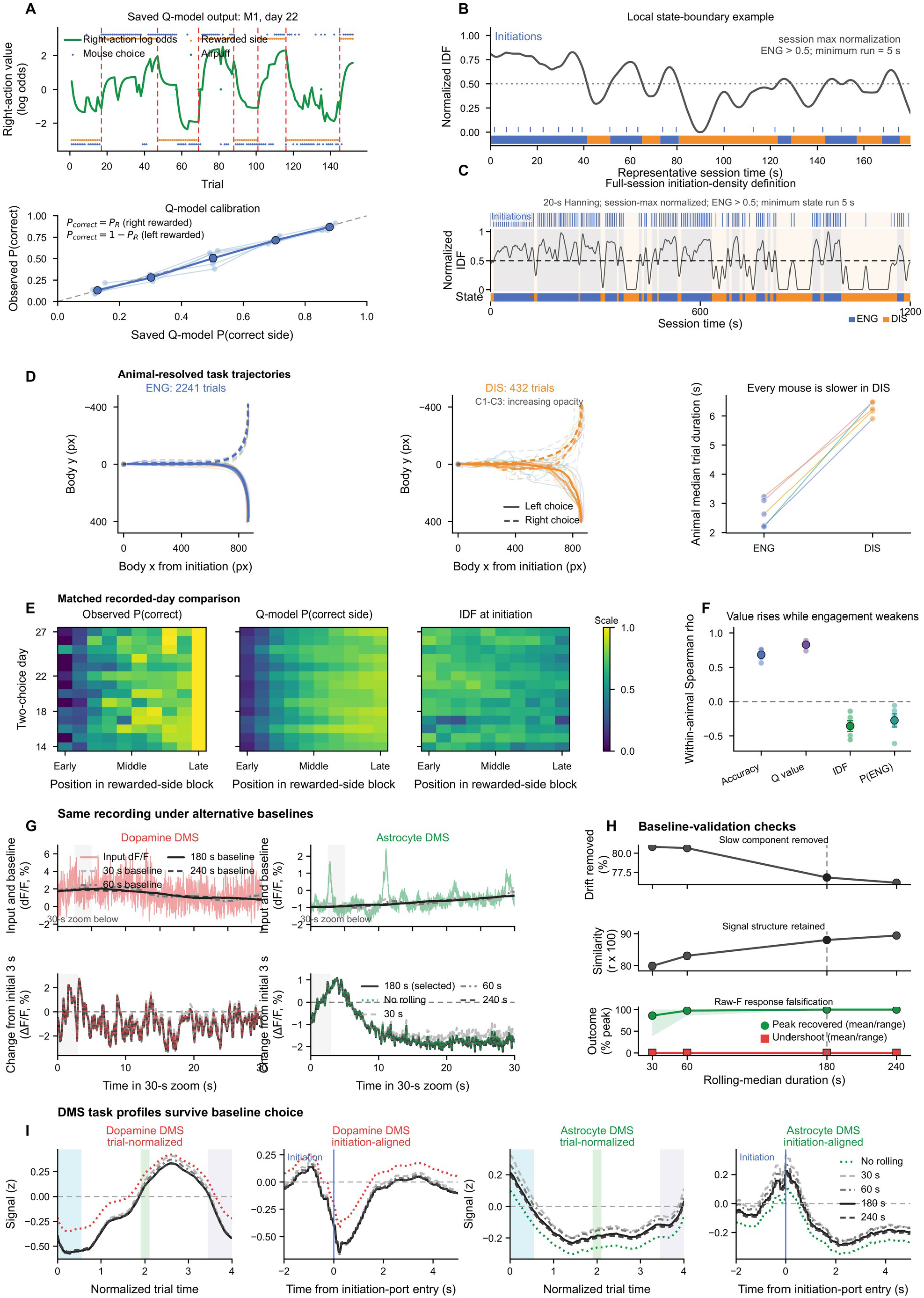
Behavioral-state definition and photometry preprocessing. (A) Example fitted action values and choice probabilities in the probabilistic two-choice task. (B and C) Initiation density (IDF), trial initiations, and ENG/DIS assignments in an expanded interval and a complete session. ENG/DIS assignments used behavior only. (D) Initiation-to-choice trajectories and movement times in ENG and DIS; thin traces and paired points are mice. (E) Accuracy, fitted action value, IDF, and ENG occupancy across normalized reward-block progression. Rows are recording days. (F) Mouse-level Spearman correlations (ρ) between block progression and each measure. (G) Example signal and rolling-median baselines, with an expanded interval below. (H) Baseline-window comparisons for drift removal, signal preservation, and recovery of constructed event peaks and undershoots. (I) Trial-normalized and initiation-aligned dopamine and astrocyte profiles across baseline choices. The selected baseline was a 180-s rolling median. These comparisons use five mice and 62 sessions, with four signal-region traces per session. Thick behavioral curves summarize mice; error bars show SEM where displayed. Analysis procedures: Methods, M10.5, M8, M10.2, and M5.3.

### 2 Temporal progression and recent dopamine history

#### State separation and temporal position

Classification in held-out sessions confirmed that stable ENG-DIS separation was present in the original feature sets, beyond the low-dimensional visualization. Balanced accuracy was 0.917 for astrocyte features, 0.925 for dopamine features, and 0.873 for features describing the relationship between the signals; all five mouse means were above chance for each family (fig. S5, A to E). The relation features combined coupling, timing, and response measurements, showing that state information was also present in how the signals interacted.

Relation features distinguished early from core phases during ENG (balanced accuracy 0.788; permutation P=0.0099); the corresponding DIS accuracy was 0.542 (P=0.0891; fig. S5F). Individual-feature analyses identified both prediction by each measurement alone and its additional information beyond the other features in its family (fig. S5G; Table S1). Phase organization persisted across window durations and embedding initializations (fig. S5, H and I; Methods, M11.2).

#### Response timing and task-event controls

Absolute regional response curves preserved the transition-related differences before subtraction of the stable-state control (fig. S5O). The centered comparison in the main text therefore reflected differences already present in the regional response profiles (Methods, M11.4).

The timing controls clarify the relationship between dopamine-event timing and the evolving astrocyte response. For dopamine events preceding C1, the early 1-6-s window could cross the C1 boundary, whereas the later 10-20-s window usually fell after it (fig. S5N). Early response gain was greatest before ENG entry, while later activity was greatest in C3 (fig. S6, B and C). Complete regional waveforms show the broader changes in response shape sampled by these windows (fig. S6, F and G; Methods, M11.1 and M11.4).

The fraction of dopamine events occurring near task actions increased through the IDF rise, and action composition differed across the cycle (fig. S5, K and L). Held-mouse comparisons identified additional astrocyte phase information beyond task-event information, particularly during ENG (fig. S5M). Aligning astrocyte activity to the same action, trial initiation, provided a separate control for changing action composition: response waveforms varied with trajectory position, and the state-by-phase contrast exceeded time-shifted controls (fig. S6, H and I). The phase dependence therefore could not be reduced to how often the sampled actions occurred.

#### Astrocyte calcium sequence and selective boundary prediction

Waveform categories (“signatures”) capture differences in preceding astrocyte activity and the early and later response shapes around dopamine events. With four, five, or six categories, the forward sequence model predicted the next category better than the reversed transition model in all five mice during both ENG and DIS (fig. S6J). The size of this advantage varied with the number of categories, but its direction was preserved (Methods, M11.5).

**Fig. S3.**
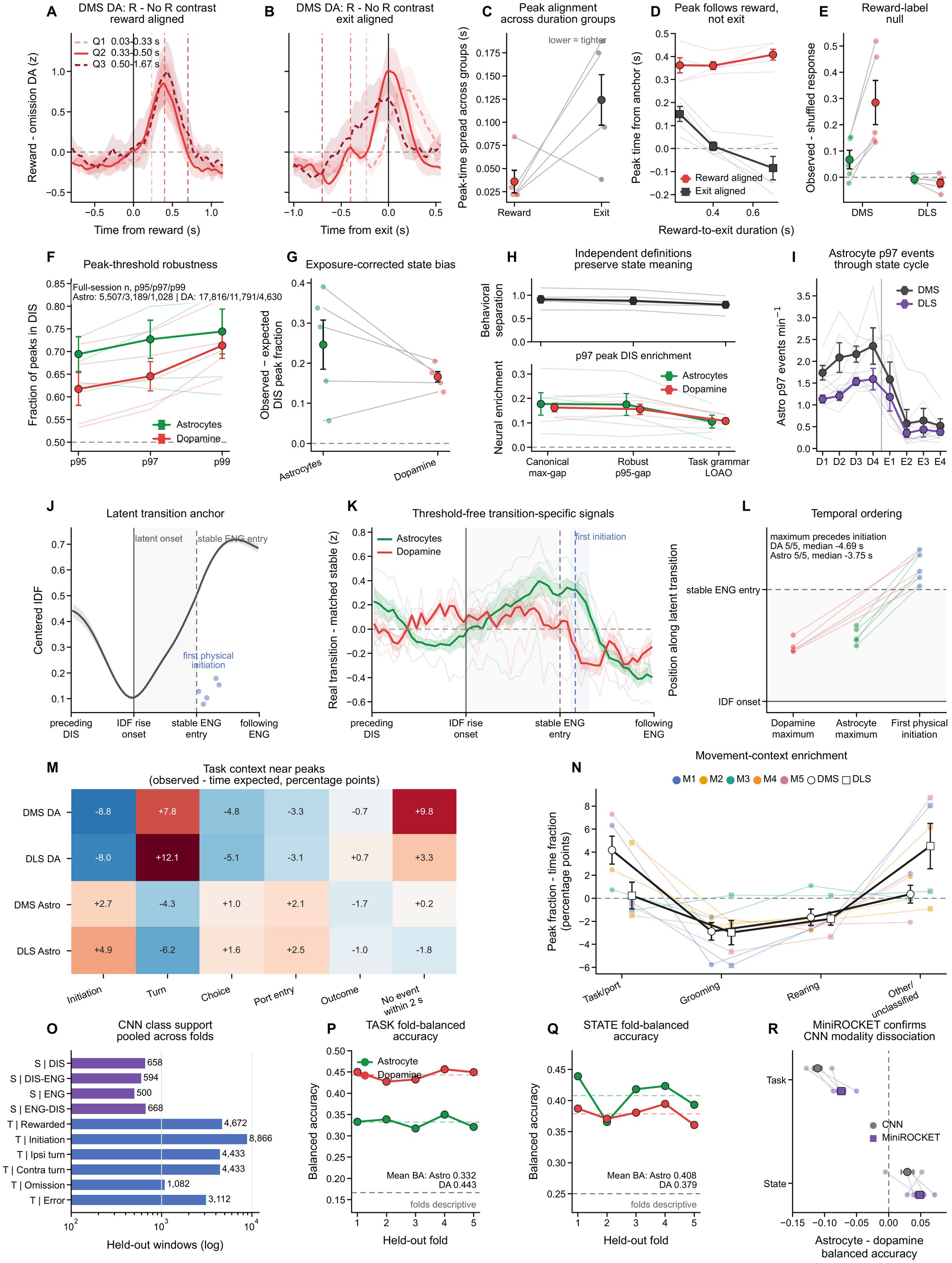
Outcome alignment, state robustness, and decoding controls. (A and B) DMS dopamine reward-minus-omission responses aligned to reward delivery or port exit, separated into three reward-to-exit duration groups. (C and D) Across-group spread of response-peak latency and its relationship to reward-to-exit duration. (E) Reward-omission contrasts relative to shuffled reward labels for dopamine and astrocyte calcium in DMS/DLS. (F and G) DIS peak fractions at the 95th-, 97th-, and 99th-percentile thresholds, before and after subtracting DIS time occupancy. (H) Behavioral separation and peak enrichment under alternative behavior-only state definitions. (I) Astrocyte peak rates across equal quarters of DIS and ENG. (J) IDF-rise alignment and the location of the first renewed initiation. (K and L) Continuous signal changes relative to matched stable DIS and their timing around re-engagement; no peak threshold is used. (M and N) Task-event and movement-motif enrichment around large peaks, calculated relative to expected time occupancy. (O) Numbers of held-out examples by class. (P and Q) Held-out-fold balanced accuracy for task-event and state decoding. (R) Astrocyte-minus-dopamine decoding differences using the InceptionTime classifier and MiniROCKET. Five mice contributed 62 sessions. Pale points or lines show mouse summaries, except the fold-level decoding displays. Positive enrichment indicates overrepresentation relative to the relevant timing or label control. Analysis procedures: Methods, M8 and M10.

Boundary prediction tested whether astrocyte activity helped anticipate entry, progression between defined phases, or exit along ENG and DIS trajectories. Specifically, the model predicted whether the next boundary would occur before the next dopamine event. Accumulated preceding activity contributed at DIS entry (+0.0586 bits per event; all five mice positive) and early progression, whereas current waveforms contributed at late ENG progression and exit (+0.0209 and +0.0150 bits per event; fig. S6, K and L). Restricting ENG-entry prediction to information available at prediction time did not yield reliable improvement (fig. S6M). Thus, astrocyte activity provided additional predictive information at particular points in the engagement cycle.

#### Contributions of preceding dopamine and astrocyte activity

Recent dopamine activity contributed to predicting astrocyte responses beyond current dopamine, movement, and behavioral phase. Adding dopamine history increased held-session R² by 0.044 for the early response and 0.070 for the later response (0.051 for the joint response; fig. S6D). Phase also added information after dopamine history, with a larger increment for the later response (ΔR²=0.018) than the early response (0.003). These comparisons distinguish the contribution of recent dopamine from the additional phase dependence of the evolving astrocyte response.

Matched feature comparisons showed that phase and progression within phase explained more variation in astrocyte context and response features than in current or recent dopamine features (fig. S6E). The complementary accumulated-dopamine model improved prediction in all five mice (exact two-sided animal-level P=0.0625). Prediction and trajectory-reconstruction procedures are described in Methods, M11.7.

The astrocyte signal’s own preceding activity also added predictive information. Adding it to dopamine, immediate behavior, and state/history predictors increased explained variance in the subsequent response from R²=0.103 to 0.237 and exceeded a timing-shift control (fig. S5J; Methods, M11.7).

**Fig. S4.**
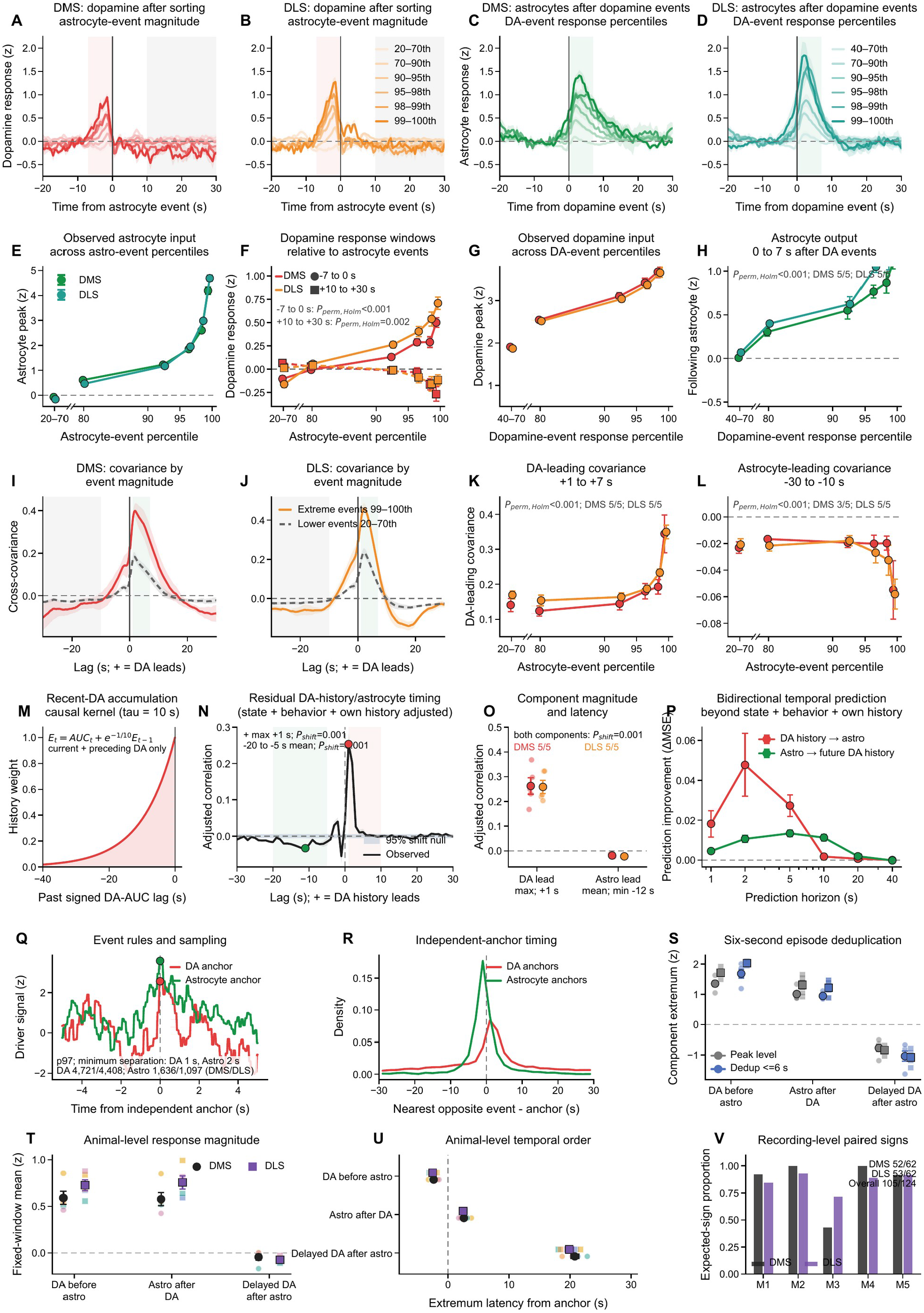
Temporal coupling between dopamine and astrocyte calcium across event magnitudes. Panels A to L describe event-centered relationships; M to P examine coupling and predictive information after accounting for behavior and preceding signal activity; Q to V examine event definitions and consistency across recordings. (A and B) Astrocyte-aligned dopamine responses in DMS and DLS, grouped by astrocyte-event magnitude. (C and D) Dopamine-aligned astrocyte responses, grouped by dopamine-event magnitude. Events were detected independently and ranked within session. (E and F) Astrocyte anchor amplitude and associated dopamine responses before (−7 to 0 s) and after (10-30 s) the astrocyte event. (G and H) Dopamine anchor amplitude and subsequent astrocyte response (0-7 s). (I and J) Event-conditioned covariance for large and lower-magnitude events. (K and L) Dopamine-leading (+1 to +7 s) and astrocyte-leading (−30 to −10 s) covariance components. (M) Signed recent-dopamine signal with a 10-s decay constant. (N) Cross-correlation after adjustment for behavior, phase, and each signal’s preceding activity; positive lag denotes dopamine leading. Shading shows the 95% shifted-null interval. (O) Mouse-level adjusted component magnitudes. (P) Reduction in held-session prediction error after adding the other signal’s preceding activity at the indicated future horizons. (Q) Independent 97th-percentile event anchors. (R) Timing of the nearest opposite-signal peak relative to each anchor. (S) Event-centered extrema using individual peaks or one peak per 6-s event cluster. (T and U) Mouse-level fixed-window responses and extremum latencies. (V) Fraction of recordings per mouse containing both the positive dopamine-leading and negative astrocyte-leading components. Both signs occurred in 52/62 DMS and 53/62 DLS recordings. Summaries show mouse mean ± SEM unless a null interval is indicated (five mice; 62 sessions). Analysis procedures: Methods, M12.

**Fig. S5.**
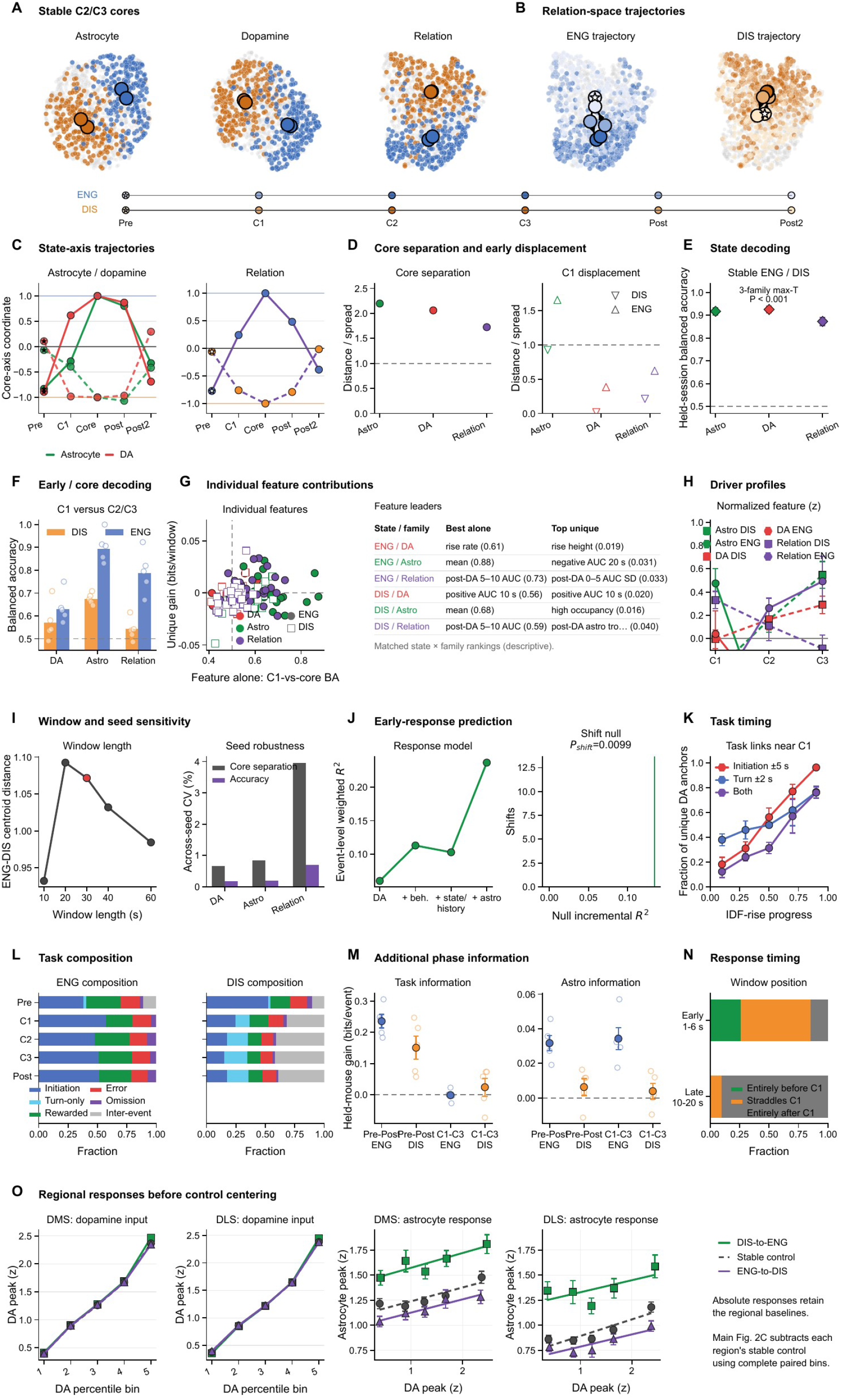
State representation and astrocyte response controls. Five mice contributed 62 sessions. (A) Stable-core UMAP representations of astrocyte, dopamine, and relation features. (B) Relation-feature trajectories through ENG and DIS. Small points are session-region-phase summaries; large symbols and arrows show phase centroids and order. (C) Phase positions along the ENG-DIS axis for astrocyte, dopamine, and relation features. (D) Stable-core separation and C1 displacement from the stable core, relative to within-state spread. (E) Held-session balanced accuracy for stable ENG versus DIS across the three feature families; chance, 0.5. (F) Animal-held-out C1-versus-C2/C3 classification within ENG and DIS. (G) Individual-feature balanced accuracy versus additional held-out information beyond the other features in the same family, with equal weight for each recording session and region; leading features are listed at right. (H) Normalized phase profiles of selected features. (I) Sensitivity to analysis-window duration and embedding seed. (J) Prediction of early astrocyte responses after successive addition of dopamine, behavior, state/history, and preceding astrocyte activity, with a timing-shift control. (K and L) Task-event timing near state entry and task composition across phases. (M) Additional phase information from task and astrocyte measurements in held-mouse comparisons. (N) Placement of early (1-6 s) and secondary (10-20 s) response windows relative to C1. In the bout-quarter framework used for A to N, C1-C3 are the first three quarters, Post is the fourth quarter, and Pre/Post2 are exterior flanks. (O) Original absolute dopamine-input and astrocyte-response curves for DMS and DLS at matched dopamine-amplitude bins, retaining all available regional measurements. Main Figure 2C uses the complete paired subset and centers astrocyte responses on the corresponding stable control. Small points or thin lines are animal or session summaries as specified in Methods; error bars show SEM. Feature definitions and contributions: table S1. Aggregation, models, and tests: Materials and Methods, M11 and M11.2.

#### Directionally asymmetric coupling and event controls

Event-magnitude analyses showed that the temporal relationship described in the main Results scaled with signal size. Larger astrocyte events were preceded by larger dopamine elevations and followed by stronger delayed dopamine suppression in both regions. Independently detected dopamine events were followed by larger astrocyte responses, with the highest-versus-lowest magnitude contrast positive in all five mice (both regions, Holm-adjusted permutation P<0.001; fig. S4, A to H). Event-conditioned covariance showed the same ordering: positive dopamine-leading and negative astrocyte-leading components grew with astrocyte-event magnitude (fig. S4, I to L).

The relationship persisted after accounting for behavior, state progression, and each signal’s own preceding activity. Adjusted coupling retained a positive dopamine-leading component and a negative astrocyte-leading component, with the same direction in all five mice in both regions (both circular-shift P=0.001; fig. S4, N and O). Matched prediction models likewise showed different timescales and signs: dopamine history most improved prediction of astrocyte activity 2 s later, whereas astrocyte history most improved prediction of dopamine accumulation 5 s later, with positive and negative net coefficients, respectively (fig. S4P). The dopamine measure retained both elevations and reductions, allowing the analysis to capture delayed suppression (fig. S4M; Methods, M12.2-M12.3).

Independent event detection and consolidation of nearby peaks preserved the dopamine-before-astrocyte and delayed-dopamine-suppression ordering (fig. S4, Q to U). Both expected whole-session coupling signs occurred together in 52 of 62 DMS and 53 of 62 DLS recordings, demonstrating that the relationship was distributed across recordings (fig. S4V; Methods, M12.4).

**Fig. S6.**
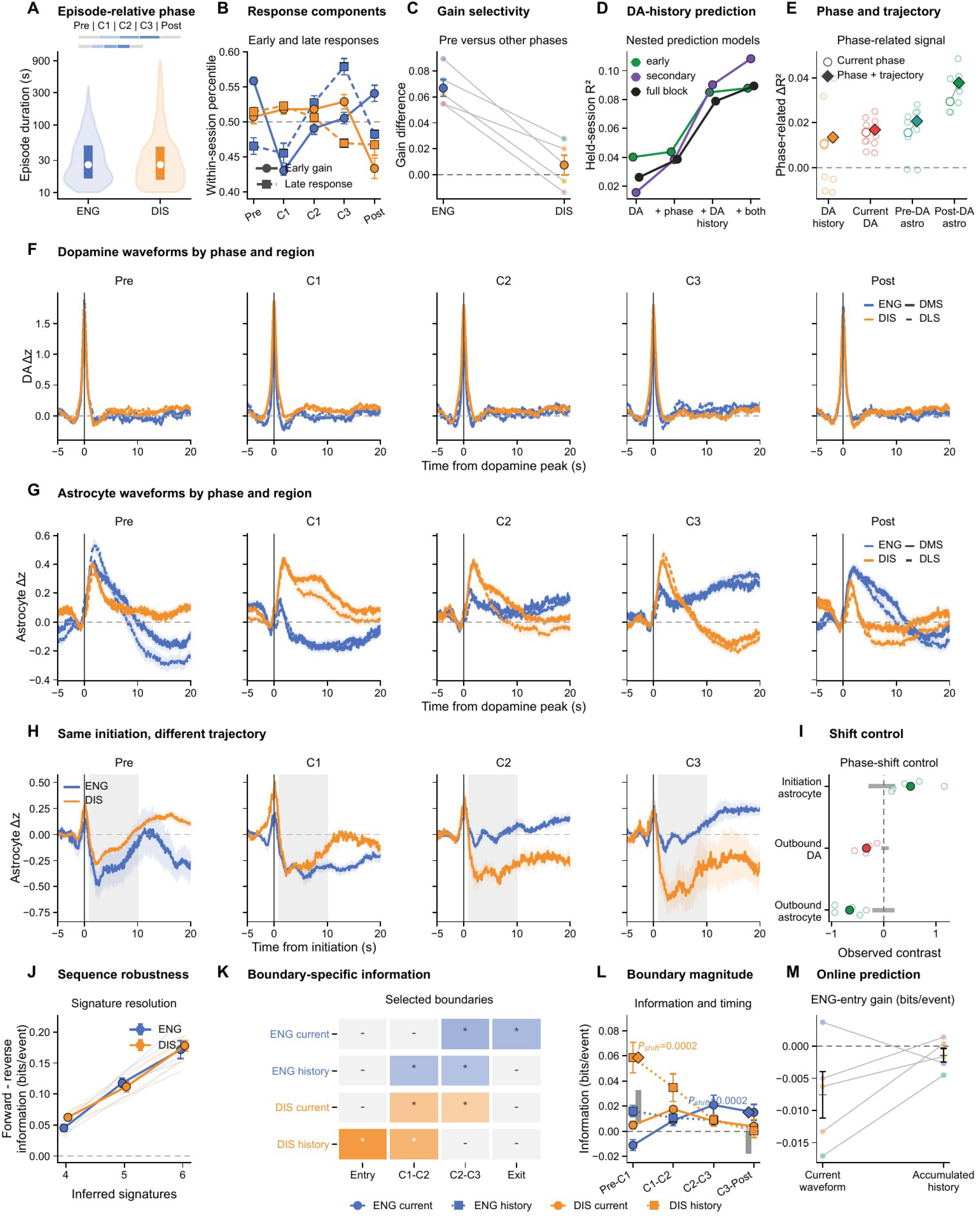
Waveform sequences, recent dopamine history, and behavioral-boundary controls. Five mice contributed 62 sessions. (A) Episode-duration distributions and equal-third phase construction: C1-C3 divide a stable episode into three equal periods, with equal-duration exterior Pre and Post flanks. (B) Early astrocyte response gain and late response across phases, expressed as within-session percentiles. (C) Early response gain before ENG entry compared with other phases. (D) Held-session prediction of early, secondary, and joint astrocyte responses after adding phase, recent dopamine history, or both to current-dopamine and movement predictors; 25,099 event-phase memberships contribute. (E) Variation explained by phase and within-phase progression in recent dopamine history, current dopamine, pre-dopamine astrocyte context, and post-dopamine astrocyte responses. (F and G) Dopamine-aligned dopamine and astrocyte waveforms across Pre, C1-C3, and Post; solid, DMS; dashed, DLS. (H) Initiation-aligned astrocyte responses across phases within ENG and DIS. (I) Observed contrasts relative to time-shifted controls: initiation-aligned astrocyte state-by-phase interaction, outbound dopamine ENG-DIS contrast, and outbound astrocyte Pre-C1 contrast (top to bottom). Gray intervals show shifted controls. (J) Forward-minus-reverse sequence information across different numbers of waveform signatures. (K) Boundaries with additional predictive information from current waveforms or accumulated preceding waveform history; asterisks indicate passage of both original tests. (L) Magnitude of additional boundary-prediction information and its timing-shift comparison. The predicted outcome is whether the next behavioral boundary occurs before the next dopamine event. (M) ENG-entry prediction restricted to information available at prediction time; no reliable improvement is detected. Blue, ENG; orange, DIS. Small points or thin lines show mouse summaries; error bars and waveform shading show SEM. Phase definitions, response features, models, and tests: Materials and Methods, M11 and M11.2. Extended results: Supplementary Text, section 2.

### 3 Dopamine-astrocyte interactions across timescales

#### Frequency dependence and receptor contributions

We varied SNc dopamine-neuron stimulation frequency from 5 to 60 Hz to examine how striatal dopamine release and astrocyte calcium responses scaled with stimulation frequency. Higher frequencies evoked larger dopamine responses and greater astrocyte calcium responses, with astrocyte responses strongest at 40-60 Hz (fig. S7B). Dopamine peaked earlier than astrocyte calcium in both regions (fig. S7A; Methods, M13).

Complete receptor-comparison time courses show the strong suppression of dopamine-evoked astrocyte calcium after Drd2 deletion alongside the maintained responses after Drd1 or Adra1a deletion (fig. S7, C and D). Extended adenosine traces document the prolonged control response and its marked suppression after astrocytic PMCA expression (fig. S7E). These traces provide the temporal context for the response-magnitude comparisons in the main Results.

NT5E fluorescence increased from MOR1-rich striosomal zones into the surrounding matrix, with higher matrix fluorescence in section-level comparisons (six sections from one mouse; fig. S7, G to I).

#### Dependence on recent stimulation

Fits to repeated-train responses characterized the strong astrocyte adaptation described in the main Results. An exponential decline was favored over a linear decline by ΔAIC>2 in all 14 astrocyte fits, compared with 7 of 16 dopamine fits across mouse, region, and stimulation frequency (fig. S7F). The astrocyte attenuation was therefore concentrated in the first few trains and was substantially stronger than the accompanying adaptation of dopamine release.

#### Movement controls during astrocyte stimulation

Open-field tracking tested for movement changes during the astrocyte-stimulation protocol. Speed, immobility, path length, path straightness, and immobility-bout duration showed no detectable changes during or after stimulation (fig. S7, J to L). Behavioral-motif occupancy was also similar before, during, and after stimulation (fig. S7M), extending the movement assessment beyond overall locomotor activity (Methods, M7.1-M7.2).

**Fig. S7.**
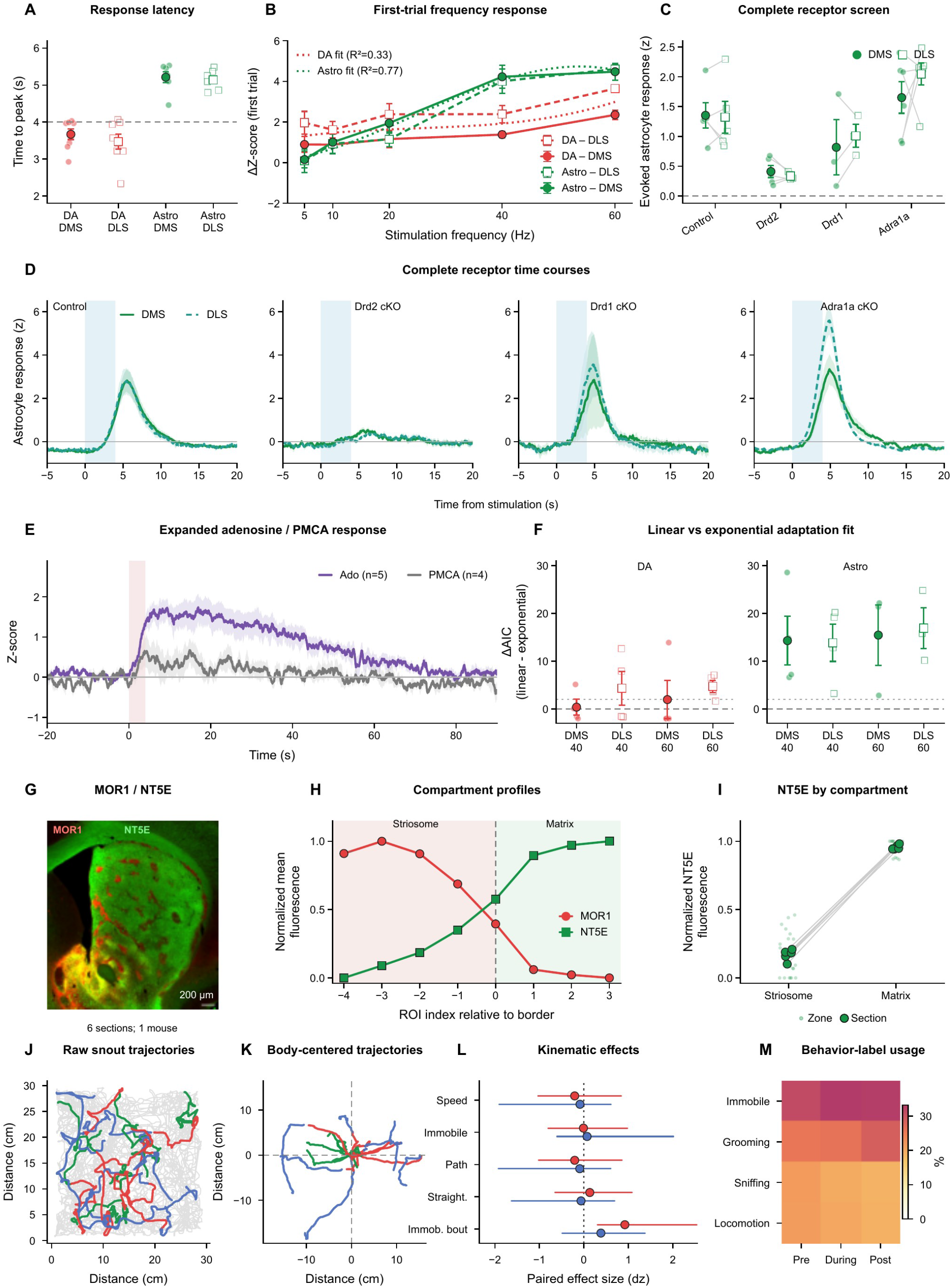
Calibration, anatomical context and controls for reciprocal dopamine-astrocyte interactions. (A) Striatal dopamine and astrocyte-calcium peak latencies after SNc dopamine-neuron stimulation; dashed line, end of the 4-s train. (B) First-train responses across stimulation frequencies. (C and D) Evoked astrocyte-calcium magnitudes and time courses in controls and after astrocytic *Drd2*, *Drd1*, or *Adra1a* deletion. Filled/solid and open/dashed symbols or traces denote DMS and DLS; paired regional measurements are connected in C. (E) Extended adenosine time courses in control (n=5 mice) and astrocyte-PMCA (n=4) cohorts. (F) Comparison of linear and exponential fits to repeated-train attenuation by region and frequency. ΔAIC is linear minus exponential AIC; values above 2 favor the exponential fit. The 14 astrocyte and 16 dopamine values are animal-by-region-by-frequency conditions, not independent animals. (G) Representative MOR1 (red) and NT5E (green) staining; scale bar, 200 µm. (H) Normalized fluorescence profiles across eight zones extending from striosomal interiors into surrounding matrix (six sections, one mouse). (I) NT5E intensity in striosome and matrix zones. Small dots show zone measurements; larger paired dots show section means weighted by zone area. The border zone is excluded from I. (J and K) Raw snout trajectories and body-centered center-of-body trajectories during dorsal-striatal astrocyte stimulation. (L) Paired effects on speed, immobility, path length, straightness, and immobility-bout duration during (red) and after (blue) stimulation relative to baseline. Points show standardized paired effects (d_z); intervals are 95% bootstrap confidence intervals. (M) Behavioral-label occupancy before, during, and after stimulation. No consistent large movement change accompanied stimulation. Small points represent subject or condition summaries as specified; response traces and other summary error bars show mean ± SEM. Cohorts and stimulation procedures: Methods, M2.1-M2.2 and M2.5. Open-field assay and movement analyses: Methods, M6.2 and M7.1-M7.2. Anatomical quantification: Methods, M3.2. Response analyses and tests: Methods, M13.

### 4 Engagement during cue-guided learning

#### Expression coverage and behavioral coordinates

Regional measurements confirmed that PMCA expression was centered in DMS and DLS, with less extensive spillover into cortex and ventral striatum (fig. S8, A and B). All seven mice had lower coverage in DLS than DMS, supporting the regional emphasis of the anatomical comparison in the main Results (Methods, M3.1).

The joint reward-cost response plots show how changes in responding to both cues distinguish overall response propensity from cue discrimination (fig. S8C). Group-balanced selectivity curves located the behavioral change between low and established discrimination (fig. S8D; Methods, M14.1-M14.2).

During the transition interval, PMCA mice shifted toward lower response probabilities for both reward and cost cues (joint permutation P=0.013; fig. S8G); the corresponding joint comparisons at low and established discrimination detected no group difference (fig. S8, F and H). The overlapping-window scan placed its largest separation in the same intermediate range (maximum-cluster-mass P=0.112; fig. S8E). This scan was distinct from the prespecified transition-versus-flanking-stages contrast reported in the main Results.

#### Matched engagement and response controls

Joint response plots at matched low, middle, and high engagement showed similar reward-cost response patterns across groups (fig. S8, I to K), supporting the main-text comparison of discrimination at matched engagement.

Licking controls separated the probability of making a response from licking once a response occurred. Response-lick distributions on trials with a pre-outcome response and consummatory licks per reward were similar between groups (P=0.9038 and 0.8838; fig. S8, L and M). Thus, reduced responding occurred with maintained licking on response trials and during reward consumption. The performance distribution also shows the lower proportion reaching high discrimination: six of eight controls and one of seven PMCA mice exceeded maximum-session d′>1.2 (P=0.0406; fig. S8N).

The same eight control and seven PMCA mice had previously performed simpler tasks testing reward earning, effort, withholding, and cue responding. Across six prespecified endpoints, PMCA mice showed no broad performance reduction (fig. S9A). They made more licks per earned reward in late fixed-ratio-1 training without a lower reward rate (exact P=0.0016; BH-adjusted q=0.028; fig. S9B). These tests extend the licking controls to the ability to obtain rewards and perform operant actions (Methods, M14.4).

**Fig. S8.**
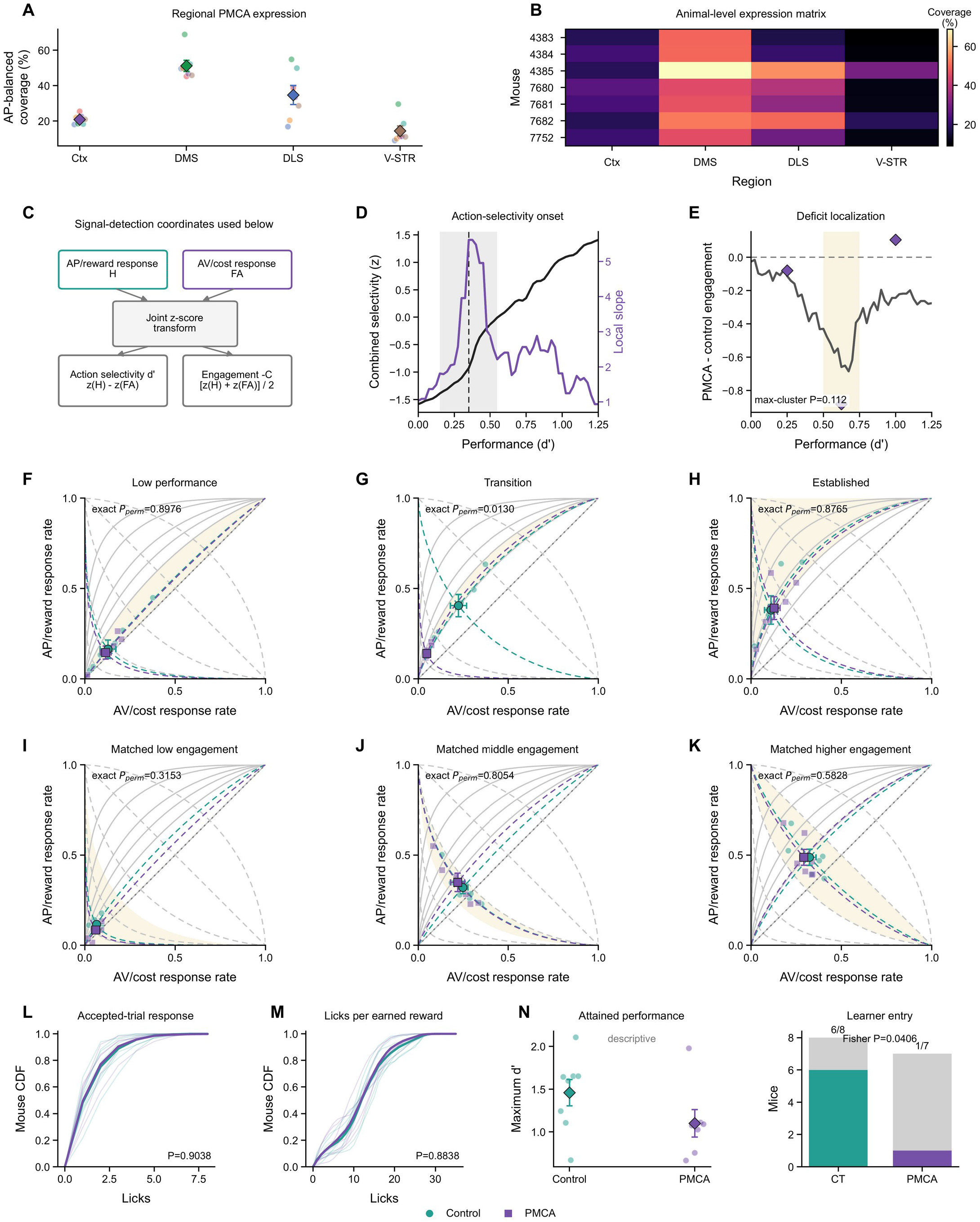
Regional PMCA expression and behavioral components of reduced engagement. Seven PMCA mice and eight GFAP-Cre controls were compared. (A and B) Regional expression coverage, averaged across anterior-posterior levels, and individual-mouse coverage in cortex, DMS, DLS, and ventral striatum. (C) Conversion of reward- and cost-cue response rates into discrimination (d′) and engagement (−C). (D) Group-balanced cue-selectivity curve and its local slope across discrimination levels. (E) PMCA-minus-control engagement across overlapping performance windows; diamonds mark the fixed low, transition, and established stages. The trajectory-wide maximum-cluster test gives P=0.112. (F to H) Reward- versus cost-cue response probabilities at each fixed performance stage. (I to K) The same response geometry at matched low, middle, and high engagement. Solid and dotted contours indicate constant discrimination and engagement; shading marks the selected bin. During transition, PMCA mice responded less to both cues; matched-engagement comparisons showed no detectable discrimination difference. (L and M) Cumulative lick-count distributions on responded trials and per earned reward. (N) Maximum-session discrimination and numbers of mice exceeding d′>1.2 (1/7 PMCA and 6/8 controls). Teal circles, controls; purple squares, PMCA. Pale points or curves are mice; large symbols and thick curves summarize groups, with SEM where shown. Expression quantification and behavioral analyses: Methods, M3.1 and M14.

**Fig. S9.**
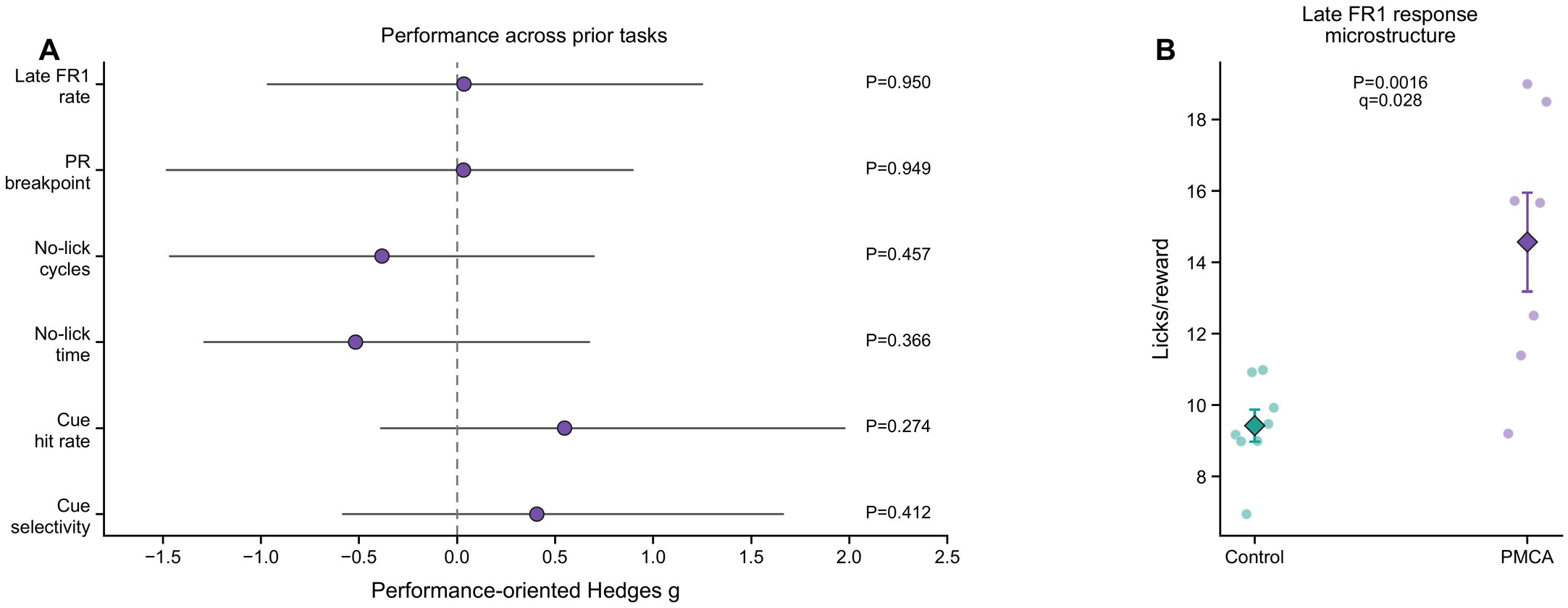
Earlier operant performance does not show a general response deficit in PMCA mice. The same seven PMCA mice and eight GFAP-Cre controls were tested before approach-avoidance learning. (A) Standardized group differences in late fixed-ratio-1 reward rate, progressive-ratio breakpoint, withholding-task cycles and completion time, and variable-interval hit rate and cue selectivity. Cue selectivity uses cue and intertrial lick rates in Hz; the cue count over the 2-s window was divided by two. Positive Hedges’ g denotes better PMCA performance; horizontal intervals are 95% mouse-bootstrap confidence intervals and zero denotes no group difference. (B) Licks per earned reward late in fixed-ratio-1 training. PMCA mice made more licks per reward despite similar reward rates. Pale points are mice; diamonds and error bars show mean ± SEM. P values use exact mouse-label permutation. The primary performance endpoints showed no broad group deficit. Analysis procedures and multiplicity correction: Methods, M14.4.

### Supplementary Table S1

*Analytical descriptions and predictive contributions of dopamine, astrocyte-calcium, and dopamine-astrocyte relation features for distinguishing early (C1) from stable-core (C2/C3) phases within ENG and DIS (fig. S5, F to I)*.

#### State-phase information carried by individual signal features

Standalone balanced accuracy (BA; 0.5 = chance) distinguishes C1 from C2/C3 within ENG or DIS using one feature, with region included as a common covariate. Unique bits/window is the held-out log-loss improvement per session-region-condition feature vector when the feature is added to the other features in its family. Each vector summarizes corresponding local windows within a recording session and region. Positive values indicate additional phase information; negative values indicate poorer held-out prediction after adding the feature. The table contains all 64 features: 17 astrocyte-calcium, 19 dopamine, and 28 dopamine-astrocyte relation measurements. Model procedures are described in Methods, M11.2.

| # | Family | ID | Feature | Analytical description | ENG BA /<br>unique bits | DIS BA /<br>unique bits |
| --- | --- | --- | --- | --- | --- | --- |
| 1 | Astro | AS06 | Astro dip duration | Mean duration of negative astrocyte-calcium excursions below the local baseline (s); longer values indicate more persistent suppression. | 0.490 / -0.001 | 0.552 / -0.004 |
| 2 | Astro | AS03 | Astro rise duration | Mean duration of positive astrocyte-calcium excursions above the local baseline (s); longer values indicate more persistent activation. | 0.597 / 0.001 | 0.581 / -0.010 |
| 3 | Astro | AS05 | Astro dip rate | Number of negative astrocyte-calcium excursions per minute; higher values indicate more frequent dips. | 0.722 / -0.004 | 0.637 / -0.003 |
| 4 | Astro | AS02 | Astro rise rate | Number of positive astrocyte-calcium excursions per minute; higher values indicate more frequent rises. | 0.593 / -0.006 | 0.593 / -0.000 |
| 5 | Astro | AS20 | Astro rise-time SD | SD of astrocyte-event onset-to-peak time (s); higher values indicate more variable rise timing. | 0.639 / -0.004 | 0.544 / -0.001 |
| 6 | Astro | AS07 | Astro dip depth | Mean depth of negative astrocyte-calcium excursions relative to the local baseline (z); higher values indicate deeper dips. | 0.677 / -0.018 | 0.645 / 0.011 |
| 7 | Astro | AS01 | Astro peaks | Mean amplitude of detected astrocyte-calcium peaks within the state window (z); higher values indicate stronger transients. | 0.669 / -0.002 | 0.520 / -0.007 |
| 8 | Astro | AS04 | Astro rise height | Mean height of positive astrocyte-calcium excursions relative to the local baseline (z); higher values indicate larger rises. | 0.712 / -0.005 | 0.474 / -0.008 |
| 9 | Astro | AS22 | Astro negative AUC 20 s | Mean area below baseline during the 20 s after an astrocyte event, relative to the preceding 20-s mean (z s). Area is expressed as a positive magnitude; larger values indicate a stronger or more sustained negative tail. | 0.641 / 0.031 | 0.458 / -0.020 |
| 10 | Astro | AS19 | Astro local-dip AUC | Mean area below the local window baseline 10-20 s after a positive astrocyte-calcium excursion (z s); higher values indicate a stronger or longer undershoot. | 0.613 / 0.016 | 0.631 / -0.015 |
| 11 | Astro | AS21 | Astro positive AUC 20 s | Mean area above baseline during the 20 s after an astrocyte event, relative to the preceding 20-s mean (z s); larger values indicate a stronger or more sustained positive tail. | 0.550 / 0.025 | 0.427 / -0.049 |
| 12 | Astro | AS17 | Astro high occupancy | Fraction of time astrocyte calcium exceeds its session-wide 85th percentile; higher values indicate more sustained high activity. | 0.825 / -0.009 | 0.675 / 0.016 |
| 13 | Astro | AS18 | Astro low occupancy | Fraction of time astrocyte calcium is below its session-wide 25th percentile; higher values indicate more sustained low activity. | 0.730 / -0.022 | 0.597 / -0.015 |
| 14 | Astro | AS15 | Astro maximum | Maximum astrocyte-calcium signal in each state window, averaged across windows (z); captures the upper extreme of activity. | 0.859 / 0.019 | 0.613 / -0.006 |
| 15 | Astro | AS14 | Astro mean | Mean astrocyte-calcium signal in each state window, averaged across windows (z); captures the sustained activity level. | 0.875 / -0.004 | 0.679 / -0.006 |
| 16 | Astro | AS16 | Astro minimum | Minimum astrocyte-calcium signal in each state window, averaged across windows (z); captures the lower extreme of activity. | 0.694 / -0.009 | 0.613 / 0.005 |
| 17 | Astro | AS23 | Astro peak timing 40 s | Mean time to the maximum astrocyte-calcium signal within 0-40 s after an astrocyte reference event (s); larger values indicate a later maximum. | 0.635 / 0.023 | 0.548 / 0.003 |
| 18 | DA | DA06 | DA dip duration | Mean duration of negative dopamine excursions below the local baseline (s); longer values indicate more persistent suppression. | 0.526 / -0.007 | 0.472 / -0.000 |
| 19 | DA | DA13 | DA long dips | Fraction of negative dopamine excursions lasting at least 3 s; higher values indicate more prolonged dips. | 0.520 / -0.004 | 0.460 / -0.006 |
| 20 | DA | DA10 | DA long rises | Fraction of positive dopamine excursions lasting at least 3 s; higher values indicate more prolonged rises. | 0.417 / -0.012 | 0.548 / -0.007 |
| 21 | DA | DA03 | DA rise duration | Mean duration of positive dopamine excursions above the local baseline (s); longer values indicate more persistent activation. | 0.504 / -0.009 | 0.554 / 0.000 |
| 22 | DA | DA11 | DA short dips | Fraction of negative dopamine excursions lasting less than 1.5 s; higher values indicate briefer dips. | 0.522 / -0.006 | 0.458 / 0.006 |
| 23 | DA | DA08 | DA short rises | Fraction of positive dopamine excursions lasting less than 1.5 s; higher values indicate briefer rises. | 0.506 / -0.005 | 0.466 / -0.002 |
| 24 | DA | DA21 | DA event width | Mean width of dopamine events at half prominence (s); higher values indicate broader transients. | 0.472 / -0.008 | 0.548 / 0.007 |
| 25 | DA | DA05 | DA dip rate | Number of negative dopamine excursions per minute; higher values indicate more frequent dips. | 0.532 / -0.005 | 0.508 / -0.005 |
| 26 | DA | DA02 | DA rise rate | Number of positive dopamine excursions per minute; higher values indicate more frequent rises. | 0.609 / 0.012 | 0.512 / 0.001 |
| 27 | DA | DA07 | DA dip depth | Mean depth of negative dopamine excursions relative to the local baseline (z); higher values indicate deeper dips. | 0.528 / -0.006 | 0.522 / -0.020 |
| 28 | DA | DA01 | DA peaks | Mean amplitude of detected dopamine peaks within the state window (z); higher values indicate stronger transients. | 0.550 / 0.005 | 0.411 / -0.009 |
| 29 | DA | DA04 | DA rise height | Mean height of positive dopamine excursions relative to the local baseline (z); higher values indicate larger rises. | 0.556 / 0.019 | 0.488 / -0.008 |
| 30 | DA | DA20 | DA negative AUC 20 s | Mean area below baseline during the 20 s after a dopamine event, relative to the preceding 20-s mean (z s). Area is expressed as a positive magnitude; larger values indicate a stronger or more sustained negative tail. | 0.571 / -0.006 | 0.450 / -0.005 |
| 31 | DA | DA19 | DA positive<br>AUC 10 s | Mean area above baseline during the 10 s after a dopamine event, relative to the preceding 10-s mean (z s); larger values indicate a stronger or more sustained positive tail. | 0.548 / 0.001 | 0.558 / 0.020 |
| 32 | DA | DA17 | DA high<br>occupancy | Fraction of time dopamine exceeds its session-wide 85th percentile; higher values indicate more sustained high activity. | 0.528 / -0.002 | 0.540 / -0.009 |
| 33 | DA | DA18 | DA low<br>occupancy | Fraction of time dopamine is below its session-wide 25th percentile; higher values indicate more sustained low activity. | 0.558 / -0.000 | 0.540 / -0.003 |
| 34 | DA | DA15 | DA maximum | Maximum dopamine signal in each state window, averaged across windows (z); captures the upper extreme of activity. | 0.609 / -0.004 | 0.464 / -0.001 |
| 35 | DA | DA14 | DA mean | Mean dopamine signal in each state window, averaged across windows (z); captures the sustained activity level. | 0.413 / -0.000 | 0.488 / -0.004 |
| 36 | DA | DA16 | DA minimum | Minimum dopamine signal in each state window, averaged across windows (z); captures the lower extreme of activity. | 0.550 / -0.008 | 0.478 / -0.002 |
| 37 | Relation | REL008 | Astro-leading<br>correlation | Mean event-aligned correlation at lags where astrocyte activity leads dopamine by 10-30 s; more negative values indicate a stronger delayed dopamine reduction after astrocyte activity. | 0.488 / -0.002 | 0.544 / -0.000 |
| 38 | Relation | REL009 | Window<br>covariance | Zero-lag covariance of dopamine and astrocyte calcium across same-state windows (z<sup>2</sup>); positive values indicate co-fluctuation in the same direction. | 0.512 / 0.012 | 0.520 / 0.009 |
| 39 | Relation | REL010 | Covariance<br>peak lag | Lag of the greatest dopamine-astrocyte covariance within $\pm 30$ s, excluding the central $\pm 1$ s (s); positive lags indicate dopamine leading astrocytes. | 0.562 / 0.004 | 0.524 / 0.001 |
| 40 | Relation | REL013 | DA-leading<br>covariance<br>minimum | Minimum covariance at lags where dopamine leads astrocytes by 1-30 s (z<sup>2</sup>); more negative values indicate stronger opposing fluctuations. | 0.567 / -0.003 | 0.512 / 0.005 |
| 41 | Relation | REL014 | DA-leading<br>minimum lag | Lag of minimum covariance on the dopamine-leading side, between 1 and 30 s (s); larger values indicate a longer dopamine lead. | 0.496 / 0.001 | 0.492 / -0.008 |
| 42 | Relation | REL011 | Astro-leading<br>covariance<br>minimum | Minimum covariance at lags where astrocytes lead dopamine by 1-30 s (z<sup>2</sup>); more negative values indicate stronger opposing fluctuations. | 0.589 / 0.012 | 0.482 / 0.001 |
| 43 | Relation | REL012 | Astro-leading<br>minimum lag | Lag of minimum covariance on the astrocyte-leading side, between -30 and -1 s (s); more negative values indicate a longer astrocyte lead. | 0.526 / -0.005 | 0.435 / -0.004 |
| 44 | Relation | REL028 | Astro/DA<br>magnitude ratio | Mean astrocyte-to-dopamine event-magnitude ratio across matched dopamine-amplitude groups. Magnitude combines positive peak, positive area, half-prominence width, and prominence as a geometric mean; higher ratios indicate stronger astrocyte recruitment per dopamine response. | 0.484 / 0.019 | 0.526 / -0.011 |
| 45 | Relation | REL029 | Astro/DA ratio IQR | Interquartile range of the astrocyte-to-dopamine event-magnitude ratio defined in REL028; higher values indicate less consistent coupling. | 0.540 / 0.008 | 0.516 / -0.012 |
| 46 | Relation | REL005 | Post-DA astro peak | Mean astrocyte-calcium maximum during 0-6 s after a dopamine event, relative to the preceding 2-s mean (z); measures the early calcium increase. | 0.534 / 0.010 | 0.476 / -0.011 |
| 47 | Relation | REL001 | Pre-astro DA peak | Mean dopamine maximum in the 15 s before an astrocyte event, relative to the mean during the 2 s after that event (z); measures preceding dopamine activity. | 0.532 / 0.006 | 0.538 / 0.004 |
| 48 | Relation | REL031 | Post-DA 0-5 AUC SD | SD of signed astrocyte-calcium area during 0-5 s after dopamine events, relative to the preceding 2-s mean (z s); higher values indicate less consistent responses. | 0.617 / 0.033 | 0.550 / -0.011 |
| 49 | Relation | REL035 | Post-DA 10-20 AUC SD | SD of signed astrocyte-calcium area during 10-20 s after dopamine events, relative to the preceding 2-s mean (z s); higher values indicate less consistent responses. | 0.637 / -0.013 | 0.498 / -0.005 |
| 50 | Relation | REL037 | Post-DA 20-30 AUC SD | SD of signed astrocyte-calcium area during 20-30 s after dopamine events, relative to the preceding 2-s mean (z s); higher values indicate less consistent responses. | 0.609 / -0.002 | 0.538 / -0.007 |
| 51 | Relation | REL033 | Post-DA 5-10 AUC SD | SD of signed astrocyte-calcium area during 5-10 s after dopamine events, relative to the preceding 2-s mean (z s); higher values indicate less consistent responses. | 0.669 / 0.005 | 0.433 / -0.017 |
| 52 | Relation | REL026 | Post-DA astro peak-lag SD | SD of time to the astrocyte-calcium maximum within 0-6 s after dopamine events (s); higher values indicate less consistent peak timing. | 0.544 / -0.004 | 0.440 / -0.010 |
| 53 | Relation | REL025 | Post-DA negative-area SD | SD of astrocyte-calcium area below the preceding 2-s baseline during 0-6 s after dopamine events (z s); higher values indicate less consistent negative responses. | 0.694 / -0.000 | 0.510 / 0.006 |
| 54 | Relation | REL024 | Post-DA astro trough SD | SD of astrocyte-calcium minima during 0-6 s after dopamine events, relative to the preceding 2-s mean (z); higher values indicate less consistent minima. | 0.639 / -0.001 | 0.562 / 0.040 |
| 55 | Relation | REL016 | Pre-astro DA response SD | SD of mean dopamine during the 6 s before astrocyte events, relative to the mean during the 2 s after each event (z); higher values indicate less consistent preceding dopamine. | 0.579 / -0.008 | 0.504 / -0.005 |
| 56 | Relation | REL003 | Post-astro DA trough | Mean dopamine minimum 10-30 s after an astrocyte event, relative to the mean from 15 to 5 s before it (z); more negative values indicate a deeper delayed dopamine trough. | 0.486 / -0.001 | 0.530 / 0.000 |
| 57 | Relation | REL030 | Post-DA 0-5 AUC | Mean signed astrocyte-calcium area during 0-5 s after dopamine events, relative to the preceding 2-s mean (z s); sign and magnitude describe response direction and persistence. | 0.653 / 0.026 | 0.442 / -0.019 |
| 58 | Relation | REL034 | Post-DA 10-20 AUC | Mean signed astrocyte-calcium area during 10-20 s after dopamine events, relative to the preceding 2-s mean (z s); sign and magnitude describe response direction and persistence. | 0.698 / -0.022 | 0.567 / -0.014 |
| 59 | Relation | REL007 | Post-DA astro trough | Mean astrocyte-calcium minimum 10-30 s after a dopamine event, relative to the mean from 15 to 5 s before it (z); more negative values indicate a deeper delayed calcium trough. | 0.595 / 0.032 | 0.506 / -0.018 |
| 60 | Relation | REL036 | Post-DA 20-30 AUC | Mean signed astrocyte-calcium area during 20-30 s after dopamine events, relative to the preceding 2-s mean (z s); sign and magnitude describe response direction and persistence. | 0.688 / 0.015 | 0.526 / -0.002 |
| 61 | Relation | REL032 | Post-DA 5-10 AUC | Mean signed astrocyte-calcium area during 5-10 s after dopamine events, relative to the preceding 2-s mean (z s); sign and magnitude describe response direction and persistence. | 0.730 / 0.012 | 0.591 / -0.002 |
| 62 | Relation | REL006 | Post-DA astro peak lag | Mean delay from a dopamine event to the astrocyte-calcium maximum within the following 6 s (s); larger values indicate slower recruitment. | 0.637 / 0.002 | 0.542 / -0.006 |
| 63 | Relation | REL002 | Pre-astro DA peak lag | Mean lead time of the dopamine maximum within the 6 s preceding an astrocyte event (s); larger values indicate a longer dopamine-to-astrocyte interval. | 0.645 / 0.003 | 0.502 / -0.008 |
| 64 | Relation | REL004 | Post-astro DA peak lag | Mean delay from an astrocyte event to the dopamine maximum within the following 6 s (s); larger values indicate a later dopamine peak. | 0.450 / -0.007 | 0.496 / 0.002 |

## Materials and Methods

### M1 Animals, ethical approval, and husbandry

For the experiments, we used the DAT-Cre knock-in mouse line (Common Name: DATIREScre, Slc6a3tm1.1(cre)Bkmn; RRID: IMSR_JAX:006660). This line was purchased from The Jackson Laboratory and maintained on a C57BL/6J background. In addition to the DAT-Cre line, we utilized wild-type C57BL/6J mice (RRID: IMSR_JAX:000664) also purchased from The Jackson Laboratory. Adra1a fl/fl mice, provided by Paul Simpson (University of California, San Francisco), were crossed with DAT-Cre mice for the astrocytic Adra1a-deletion experiments.

All procedures were approved by the Committee on Animal Care at the Massachusetts Institute of Technology, which is AAALAC accredited. Both female and male experimental mice were housed under a standard 12/12-hour light/dark cycle with free access to food and water. For the T-maze probabilistic two-choice switching task, mice were single-housed and placed on water restriction during behavioral training and recordings (1.5-2.5 g hydrogel per day maintained at minimum 85% of their free-water body weight).

#### M1.1 Experimental cohorts and biological sampling units

Five mice contributed 62 repeated sessions with simultaneous dopamine and astrocyte recordings in DMS and DLS for the two-choice and state analyses. Sessions, trials, state episodes, transitions, and detected events were treated as nested observations. For the approach-avoidance experiment, the behavioral dataset comprised seven astrocyte-PMCA mice and eight control mice that received GFAP-Cre virus injections; PMCA assignment was between animals, and the same seven PMCA mice contributed expression measurements and behavior. Mouse was the independent biological sampling unit for between-group inference.

### M2 Viral injections, fiber implantation, and optogenetic stimulation

General Procedure: Mice were anesthetized with 2% isoflurane and positioned in a stereotaxic frame (Stoelting). Prior to the first incision, Buprenorphine (0.1 mg/kg) and the local analgesic Xylocaine/Lidocaine (4 mg/kg) were administered subcutaneously for pain relief. Body temperature was maintained at 36°C using a feedback-controlled heating pad. Viral injections were performed using a micropipette attached to a Quintessential Stereotaxic Injector (Stoelting). The pipette remained in place for 5 min post-injection to ensure proper viral diffusion before being slowly withdrawn. Meloxicam (5 mg/kg, subcutaneously) was administered before surgery and on postoperative days 1 and 2. A single dose of sustained-release buprenorphine (0.5 mg/kg, subcutaneously) was administered immediately after surgery. Viral constructs, suppliers, catalog identifiers, and originating publications are listed in Table M1.

#### M2.1 Dopamine-neuron stimulation with striatal dopamine or astrocyte calcium photometry

In DAT-Cre mice, AAV5-Syn-FLEX-rc[ChrimsonR-tdTomato] (Addgene #62723) (*39*) (400 nl) was injected into the SNpc (AP: −3.08 mm, ML: ±1.25 mm, DV: −4.0 mm). pAAV-hSyn-DA3h-WPRE-hGH (green-shifted DA3h; BrainVTA #PT-4721) (*40*) or AAV5-gfaABC1D-cyto-GCaMP6f (Addgene #52925) (*41*) (400 nl) were injected into the right DLS (AP: 1.1 mm, ML: 2.15 mm, DV: 1.95 mm) and DMS (AP: 1.1 mm, ML: 1.1 mm, DV: 2.2 mm). Optical recording fibers were implanted above the striatal injection sites at angles of −10° for DMS and +10° for DLS; stimulation light was delivered over dopamine-neuron cell bodies in SNc. Dopamine neurons were stimulated with 4-s trains delivered via a 593-nm laser.

#### M2.2 Astrocyte stimulation with striatal dopamine or astrocyte calcium photometry

For astrocytic stimulation, AAV5-GFAP-Cre (Addgene #105550-AAV5) and pAAV-nEF-Con/Foff 2.0-ChRmine-oScarlet (AAV8; Addgene #137161-AAV8) (*42, 43*) (total volume 400 nl) were injected into the dorsal striatum (AP: 0.86 mm, ML: 1.75 mm, DV: 2.25-1.75 mm; 0° injection angle). pAAV-hSyn-DA3h-WPRE-hGH (green-shifted DA3h; BrainVTA #PT-4721) (*40*) or AAV5-gfaABC1D-cyto-GCaMP6f (Addgene #52925) (*41*) (400 nl) was co-injected into the same dorsal-striatal target in separate dopamine- or calcium-recording cohorts. Both fibers were angled at 15°, one inward and the other outward, to target the same injected striatal volume; one delivered stimulation light, and the other collected fluorescence for photometric recording. Astrocytes were stimulated with continuous 593-nm light for 10 s (5 mW), with 10 trials and randomized inter-trial intervals of 1.5 min.

#### M2.3 Simultaneous extracellular dopamine and astrocyte calcium recordings

For simultaneous extracellular dopamine and astrocyte calcium recordings, AAV5-gfaABC1D-cyto-GCaMP6f (Addgene #52925) (*41*) and AAV2/9-hSyn-GRAB-rDA3h (BrainVTA #PT-5198) (*40*) (400 nl) were injected into the right DLS (AP: 1.1 mm, ML: 2.15 mm, DV: 1.95 mm) and left DMS (AP: 1.1 mm, ML: 1.1 mm, DV: 2.2 mm). Optical fibers were implanted above the respective injection sites. This configuration sampled both regions while avoiding two implants in the same hemisphere. Fluorescence was collected from the local tissue volume beneath each fiber tip.

#### M2.4 Astrocyte PMCA manipulation

For the approach-avoidance cohort, dorsal-striatal astrocytes were targeted bilaterally with AAV5-GfaABC1D-mCherry-hPMCA2w/b (Addgene #111568-AAV5) to attenuate astrocytic Ca2+ signaling (*21*). Control mice in this behavioral cohort received AAV5-GFAP-Cre injections (Addgene #105550-AAV5). Expression coverage was quantified as described in M3.1.

#### M2.5 Astrocytic receptor deletion and adenosine recordings

For astrocytic Drd1 or Drd2 deletion, AAV5-CMV-FLEX-SaCas9-U6-sgDrd1 or AAV5-CMV-FLEX-SaCas9-U6-sgDrd2 (Addgene plasmids #159917 and #159918, respectively; Larry Zweifel laboratory) was injected into DMS and DLS in DAT-Cre mice together with AAV5-GFAP-Cre (Addgene #105550-AAV5) and AAV5-gfaABC1D-cyto-GCaMP6f (Addgene #52925). Astrocytic Adra1a deletion was induced by injecting the same GFAP-Cre and GCaMP6f vectors into DMS and DLS in Adra1a fl/fl × DAT-Cre mice.

Striatal adenosine was recorded with rAAV-hSyn-Ado_B10 (Ado1.0)-WPRE-hGH polyA (AAV2/8; BrainVTA #PT-1348). The astrocyte-PMCA cohort additionally expressed the GfaABC1D-mCherry-hPMCA2w/b construct described in M2.4.

### M3 Histology and anatomical quantification

Mice were anesthetized with Euthasol (pentobarbital sodium and phenytoin sodium, Virbac AH Inc.) and transcardially perfused with 0.9% saline, followed by 4% (wt/vol) paraformaldehyde in 0.1 M NaKPO4 buffer (PB). Brains were dissected, post-fixed for 90 min, and stored in 20% (vol/vol) glycerin sinking solution overnight or longer. The brains were then sectioned transversely at 30 µm on a freezing microtome and stored in 0.1% sodium azide in 0.1 M PB until use. Free-floating sections were rinsed three times (2 min each) in 0.01 M NaKPO4 buffer with sodium/potassium saline and 0.2% Triton X-100 (PBS-Tx), then blocked for 20 min using TSA Blocking Reagent (PerkinElmer). Sections were incubated overnight at 4°C on a shaker with primary antibodies suspended in TSA Blocking Reagent. Following primary incubation, sections were rinsed three times in PBS-Tx and incubated with Alexa Fluor secondary antibodies (Thermo Fisher Scientific) for 2 hr at room temperature. After secondary incubation, sections were rinsed three times in 0.1 M PB. Images were processed and analyzed using <u>Fiji</u> software. Background subtraction and contrast adjustments were performed for presentation purposes, and figures were prepared using Adobe Photoshop and Illustrator.

Assessment of optical fiber placement was based on visible lesions from the fiber in the tissue. Mice with misplaced fibers or insufficient viral expression were excluded from the study.

#### M3.1 Regional expression coverage

RFP-positive area and anatomical region area were measured at nine matched anterior-posterior levels (+1.7 to−0.5 mm) in each PMCA mouse. At each level, expression coverage was the RFP-positive area divided by the regional area, combining both hemispheres before calculation. Coverage across the sampled range was summarized for each mouse by dividing the trapezoidal area under the coverage curve by the anterior-posterior span. DMS and DLS were analyzed separately; cortical and ventral-striatal measurements assessed expression outside these targets.

#### M3.2 Striosome and matrix fluorescence

MOR1 staining defined striosomal outlines in six sections from one mouse. The ImageJ analysis measured mean MOR1 and NT5E fluorescence across eight zones extending from striosomal interiors into surrounding matrix, combining outlines within each section. Section profiles were averaged and min-max normalized separately for each marker. For striosome-matrix comparisons, each section’s NT5E profile was min-max normalized before calculating area-weighted means for the four inner zones and three matrix zones; the border zone was excluded.

### M4 Two-color fiber photometry and synchronization

For simultaneous task recordings, astrocyte Ca++ activity and extracellular dopamine were recorded using GCaMP6f and the red-shifted dopamine sensor rDA3h, respectively, with a two-color fiber photometry system (Doric Lenses). This setup enabled sequential fluorescence recording at a sampling rate of 10 Hz across three channels: a reference channel (400-410 nm), a green channel (460-490 nm), and a red channel (555-570 nm). Video recordings of mouse behavior from a FLIR-Oryx 10GigE camera at 30 frames/s were synchronized with photometry signals, tracking the xy-coordinates of five body parts (tip of the tail, base of the tail, center of the body, neck, and snout/head) using <u>DeepLabCut</u> software (*44*). Task events, optogenetic stimulation, and both video and photometry frames were timestamped using <u>Bonsai</u> software (*45*), which interfaced with external hardware via an Arduino microcontroller running the Firmata-plus protocol (https://github.com/firmata/protocol).

### M5 Photometry preprocessing

#### M5.1 Reference correction

For the two-choice recordings, reference-corrected ΔF/F was calculated from raw Doric HDF5 files using linear least-squares reference fitting, as described previously (*16, 46*). For each recording site, the reference trace was fitted separately to the astrocyte-calcium and dopamine channels. The fitted reference was subtracted from each active-channel signal, and the difference was divided by the fitted reference to obtain ΔF/F.

#### M5.2 Rolling-baseline subtraction and normalization

To remove slow baseline fluctuations, we subtracted a centered rolling median from the reference-corrected ΔF/F trace and standardized the residual over the recording session. Candidate baseline durations were 30, 60, 120, 180, and 240 s at the native 10-Hz sampling rate; the selected duration was 180 s. Task and state decoders and the 64-feature matrix used baseline-normalized fluorescence. Each analysis below specifies whether measurements used these signals or the signals after 180-s rolling-median subtraction.

#### M5.3 Evaluation of baseline subtraction

We evaluated baseline subtraction by measuring drift removal, similarity to the original trace, and recovery of constructed events. Drift and Pearson correlation were calculated across 248 session-channel traces for baseline durations of 30, 60, 180, and 240 s. Constructed Gaussian events combined amplitudes of 10, 25, 50, 100, and 200 arbitrary units with six full widths at half maximum spanning 1.04-17.82 s, on a baseline of 1000 arbitrary units. Peak recovery and artificial post-response undershoot were measured after each candidate rolling median. For the 30-s example traces, ΔF/F was centered on the median of the first 3 s and retained in its original units.

Task-response profiles were compared using the same events for every baseline duration. Signals were interpolated piecewise between initiation, turn, and choice to align the trial sequence. Initiation-port entries were also aligned in physical time from −2 to +5 s in 50-ms bins. Session curves were averaged across the 62 recordings.

### M6 Behavioral paradigms

#### M6.1 Probabilistic two-choice switching task

We used a modified version of the probabilistic two-choice switching task (*15*), adapted to a T-maze as described previously (*16*). Water restricted mice were trained to choose between two options, with reinforcement outcomes that intermittently switched without warning. A 3-μl sucrose reward was delivered at the initiation port, while an air puff (5% of incorrect trials) served as negative reinforcement. This setup encouraged a "win-stay, lose-switch" strategy, requiring mice to adapt their choices based on changing reinforcement outcomes. One port had a 75% chance of providing a sucrose reward, and the other was non-reinforced. The reinforcement schedule switched randomly every 7-24 rewarded trials. Each behavioral session lasted 20 minutes. Video data, processed with Bonsai software, enabled real-time detection of corridor end zone entries. Two Arduino microcontrollers automated the waterspouts and interfaced with Bonsai using the Firmata protocol. Behavioral video and photometry were synchronized as described in M4.

Two-choice days 1 and 2 were an introductory stage in which selection of the correct port was followed by reward with 100% probability. The main two-choice task began on day 3, when correct choices were rewarded with 75% probability. Behavior-only sessions covered days 1 to 13 and simultaneous photometry sessions covered days 14 to 27. Session performance was the fraction of trials in which the animal selected the currently correct side, computed from correct- and incorrect-choice counters rather than reward delivery. Training-history summaries included introductory days 1 and 2; rewarded-side block, calibration, and Q-model analyses were restricted to the main probabilistic stage.

#### M6.2 Open-field assay during astrocyte stimulation

For the astrocyte-stimulation experiments, mice underwent overnight water restriction and freely explored a 30 × 30 cm arena for 15 minutes to assess locomotor activity. Astrocytes were stimulated using the protocol described in M2.2. The water restriction was used to enhance exploratory drive and general motivational state, thereby allowing evaluation of both increases and decreases in locomotor activity, and to ensure an alertness level comparable to that of mice performing the reinforcement-guided decision-making task. The arena’s transparent acrylic floor allowed for video capture from below using a high-resolution Oryx 10GigE camera (Flir) recording at 30 Hz. The video was synchronized with fiber photometry recordings of striatal astrocyte calcium or extracellular dopamine during astrocyte stimulation, enabling comparison of locomotor behavior with the corresponding signal dynamics. Kinematic and behavioral-motif analyses are described in M7.1 and M7.2.

#### M6.3 Auditory approach-avoidance discrimination

Each daily behavioral session consisted of up to 75 trials of a head-fixed approach-avoidance task with randomly interleaved reward and cost trials. Trial types were determined pseudorandomly in advance. Each trial comprised four sequential epochs: (1) a 2-s auditory cue, (2) a 1-s response period, (3) an outcome period, and (4) an inter-trial interval (ITI; 6-14 s, uniformly randomized). Distinct auditory cues for reward and cost trials were delivered via an Arduino-controlled soundboard. During the response period, a lick spout was advanced into reach of the animal, and licks were detected via an infrared beam-break lickometer. The total number of licks emitted during this 1-s response window determined the magnitude of the upcoming outcome. Licks outside this response period did not influence outcome magnitude. Following a brief (<5 ms) delay after the response period, outcomes were delivered according to trial type. On reward trials, sucrose was delivered as a series of discrete drops (4 μl each), with the total number of drops determined by response magnitude (1, 3, or 6 drops for <3, 3-4, or ≥5 licks, respectively). Reward delivery was temporally extended: individual drops were dispensed sequentially, contingent on ongoing licking, within a consumption period. On cost trials, an aversive light stimulus was delivered via an LED array for a fixed duration (10 s). Light intensity scaled with response magnitude (low, medium, high), corresponding to increasing PWM output levels calibrated to approximately 4-400 lux.

Learning criteria. Learning was defined as performance exceeding d′ = 0.7 in at least 2 of 3 consecutive sessions. Training was terminated once performance stabilized, defined as two consecutive sessions differing by <0.4 d′ and both exceeding the learning threshold.

### M7 Behavioral tracking, kinematics, and motif analysis

#### M7.1 Open-field kinematics during astrocyte stimulation

Movement was analyzed using DeepLabCut (*44*), with 1,000 frames manually labeled to identify key body parts (tip of the tail, base of the tail, center of the body, neck, and snout). Video coordinates in pixels were converted to centimeters using the arena’s dimensions.

Snout trajectories were analyzed in arena coordinates. Center-of-body trajectories were also translated to the final pre-stimulation center-of-body position and rotated using the pre-onset neck-to-snout direction.

Kinematic endpoints during astrocyte stimulation were calculated from filtered DeepLabCut center-of-body coordinates at 30 Hz, using 0.0323 cm per pixel and a centered 0.5-s moving mean for speed. Immobility was defined as speed below 2 cm/s and mobility as speed at or above 2 cm/s. Pre-stimulation, stimulation, and post-stimulation windows were −10 to 0 s, 0 to 10 s, and 10 to 20 s relative to light onset, respectively. Trial values were averaged within each of seven mice before paired two-sided Wilcoxon comparisons; the Friedman test summarized the three-period comparison. Confidence intervals for paired Cohen’s dz and mean changes were estimated using 5,000 bootstrap resamples of animals.

#### M7.2 Behavioral motif classification

Behavioral motifs were identified using <u>B-SOiD</u> (*47*), an unsupervised behavioral clustering tool that segments behavior into short movement motifs from DeepLabCut coordinates. Data from each mouse were individually processed to identify motifs. For the two-choice task, motifs were assigned to the task or port, grooming, rearing, and other or unclassified categories used in the event controls. For the open-field astrocyte-stimulation experiments, B-SOiD labels were grouped into immobility, grooming, sniffing, and locomotion, and occupancy was compared before, during, and after stimulation. Motif occupancy, movement endpoints, and their statistical comparisons were summarized within animal. Event-centered windows and exposure correction are described in M10.7. Two-choice behavior was also annotated using A-SOiD (*48*). The task-event and stimulation controls used the B-SOiD classifications.

### M8 Initiation density and engagement-state definition

#### M8.1 Initiation density

Engagement states were defined from trial-initiation timing independently of the recorded fluorescence signals. Initiations were detected from falling edges of the initiation TTL signal at 30 Hz, and their event train was smoothed with a centered 20-s Hanning window to obtain the initiation density function (IDF). The smoothed rate was divided by its session maximum; values above 0.5 defined ENG and values below 0.5 defined DIS. Runs shorter than 5 s were reassigned to the other state using their original boundaries.

#### M8.2 Alternative behavioral state definitions

Sensitivity analyses used two additional IDF definitions. One normalized initiation density to its within-session 95th percentile instead of its maximum. The other classified a centered 20-s interval as ENG when it contained at least two initiations. For the IDF sensitivity masks, ENG runs shorter than 5 s were removed before resegmenting and filling DIS runs shorter than 5 s.

A complementary state model used the organization of task actions. In 5-s bins, we measured active-choice rate, rapid initiation-to-choice rate, rapid choice-to-initiation rate, trial occupancy, and the negatives of inactive-choice rate and prolonged-port-inactivity fraction. These measures were smoothed over five bins with a Hanning window and robustly scaled. A two-component Gaussian mixture with diagonal covariance described the two patterns of task activity. Empirical transition probabilities and forward-backward probabilities were used with Viterbi decoding to assign a persistent state sequence; runs shorter than 10 s were reassigned. The component with the higher mean task-action score was designated ENG.

#### M8.3 Behavioral validation of state assignments

Behavioral validation used the task-action model fitted across the recording set. Initiation rate, correct-choice rate, median speed, and immobility were excluded from model fitting and used to characterize the assigned states. Standardized ENG-minus-DIS differences were calculated within session and averaged within mouse. A composite score averaged these four differences after reversing the sign of immobility so that positive values consistently indicated greater engagement. For validation against signal-event enrichment, scaling, mixture, and transition parameters were fitted to four mice and applied unchanged to the held-out mouse.

Body-center trajectories were aligned to initiation and followed through the subsequent choice. Trajectories were averaged first within animal, separately for ENG and DIS, leftward and rightward choices, and stable-state phase. C1, C2, and C3 denote the first, middle, and final equal-duration phases of the stable-state core. Group trajectories were calculated with equal weight for each of the five animal means. Trial duration was the elapsed time from initiation to choice and was summarized as a median within each animal and state.

### M9 Alignment of signals to task events

Session-standardized ΔF/F signals from rDA3h and GCaMP6f were aligned to trial start, initiation-port exit, turn, choice-port entry, and choice-port exit. For each event, the analysis window extended halfway toward the preceding and following events, using the median inter-event intervals across trials. The windows were concatenated to represent the trial sequence while accommodating variability in event timing.

### M10 Task-event and engagement-state analyses

Task-event and engagement-state analyses used the simultaneous-recording cohort described in M1.1. Temporal associations were tested by circularly shifting the four simultaneously recorded signals together within session; reward-outcome comparisons permuted reward labels within session and choice strata. These procedures preserved the pairing of signals and regions. Randomization statistics weighted animals equally, and exact two-sided Wilcoxon tests were also applied to the five mouse-level effects.

State enrichment measured whether signal events occurred more often in a state than expected from its duration. Local peaks were detected independently within each signal and region, primarily above the session’s 97th percentile, with 95th- and 99th-percentile thresholds used for sensitivity analyses. Exposure correction subtracted the analyzed fraction of time in DIS from the fraction of peaks in DIS. Peak fractions were averaged across regions within mouse. State enrichment and rise-versus-decline comparisons used 10,000 circular shifts within each session’s stable-state analysis window. One common offset was applied to the four peak trains while behavioral boundaries remained fixed, preserving peak counts, inter-peak intervals, and cross-signal timing. The minimum displacement was 60 s, exceeding the 20-s IDF smoothing window. Session values were averaged within mouse and region before equally weighting regions and mice; centered statistics were evaluated two-sided. Additional rate profiles divided state episodes into four equal-duration quarters.

#### M10.1 Timing of signal peaks within disengagement

To measure where peaks occurred during disengagement, we selected complete DIS bouts bounded by ENG on both sides and assigned each peak above the session’s 97th percentile to the first, middle, or final equal-duration third. Each signal-region distribution was normalized by its total eligible DIS peak count; bouts without peaks contributed no locations. Probabilities were averaged equally across regions within session, sessions within mouse, and mice.

The omnibus statistic summed the squared deviations of the three mean probabilities from 1/3. A within-bout permutation test reassigned the three third labels with a common permutation across paired signals and regions, retaining peak counts, denominators, and weights. We used 100,000 permutations and a plus-one correction. Separate animal-level tests enumerated all 6^n permutations of three labels across n mice. Pairwise within-bout comparisons swapped the two tested labels, held the third fixed, and used 100,000 permutations with Holm correction across the three pairs for each signal.

#### M10.2 Rewarded-side block analysis

Rewarded-side blocks were consecutive trials governed by the same correct side. Trial ordinal position was normalized within block and assigned to 10 bins. Correct choice, correct-side probability from the Q-learning model in M10.5, IDF at initiation, and binary ENG state were averaged within animal, day, and bin for recorded days 14 to 27. Spearman correlations with normalized block position were calculated within animal; exact two-sided Wilcoxon signed-rank tests used animal as the inferential unit.

#### M10.3 Reward timing and port departure

We compared alignment to reward delivery and port departure by dividing trials into three nonoverlapping reward-to-exit duration groups defined for the dataset. Within each mouse and duration group, the response on trials with reward omission after a correct choice was subtracted from the rewarded-trial response. Session curves were averaged within mouse. Reward alignment used choice-entry times offset by the median reward timestamp; exit alignment used the first observed port exit. Contrast curves were smoothed with a centered five-sample mean, and delayed positive peaks were selected within −0.2 to +0.8 s of reward or −0.85 to +0.25 s of exit. Peak-time spread and slopes against the group-specific median reward-to-exit duration were compared within mouse using exact Wilcoxon signed-rank tests. A reward-label control used 1,000 permutations of reward and omission labels within session, switch-count block, and physical choice side, jointly for all four signals and with equal mouse weighting.

#### M10.4 Decoding task events and engagement states

The task decoder classified initiation, ipsilateral turn, contralateral turn, correct rewarded choice, correct omission, and error from the original baseline-normalized signal windows. Relative turn labels refer to the physical recording hemisphere: the two-choice recordings sampled right DLS and left DMS. The state decoder classified stable DIS, DIS-to-ENG, stable ENG, and ENG-to-DIS. For the selected state definition, transition windows were 10 s segments beginning at the detected IDF rise or decline onset; stable windows were 10 s segments centered on the C2 midpoint.

Five outer folds were used for testing, with a grouped validation split within each outer training set. Task-decoding inputs comprised 26,598 signal windows representing 13,299 physical events at two hemispheres, with 90-frame windows spanning 1.5 s before to 1.5 s after each anchor. Partially overlapping event windows, mainly between turns and neighboring task events, were kept in the same partition. Sensitivity to window grouping was also assessed. State-decoding inputs comprised 2,420 windows of 300 frames, assigned to five fixed folds.

Convolutional-neural-network decoding used the standard <u>tsai</u> InceptionTime implementation (*49, 50*) with 50 epochs and model selection by balanced accuracy on the inner validation set. For task decoding, this score was computed over the complete validation set; for state decoding, it was the mean of validation-minibatch balanced accuracies. Learning rate, batch size, weight decay, and label smoothing were selected separately for each signal and decoding task in model sweeps. The selected configuration was held fixed across all five outer folds; full training parameters are documented in the analysis code. Loss weights were inversely proportional to training-class counts, and training used a one-cycle learning-rate schedule.

Balanced accuracy was the unweighted mean of class recalls. Paired test-window class-accuracy contrasts used McNemar tests with Benjamini-Hochberg FDR correction within each decoder. Balanced accuracy was summarized across the five held-out folds for comparative decoding.

Decoder robustness was assessed by fitting MiniROCKET (*50, 51*) separately within each outer training fold. MiniROCKET generated 9,996 fixed convolutional features per trace; a class-balanced ridge classifier was fit with regularization selected in three grouped inner folds. The transform and scaler were fitted once on the outer training set and held fixed during inner-fold ridge selection. It used the same preprocessing, class definitions, held-out observations, and outer-fold assignments as the CNN (fig. S3R).

#### M10.5 Action-value estimation using Q-learning

We calculated action values for each mouse using a Q-learning model (*52*). Completed trials, choice side, reward delivery, and air-puff delivery were identified from synchronized task events. Left- and right-port values were initialized to zero at the start of each recording. After each completed choice, the value of the chosen port was updated according to:

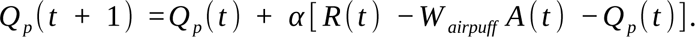

Here, *Q_p_* (*t*) is the value of port p before trial t, α is the learning rate, R(t) indicates reward delivery (1 for reward and 0 otherwise), A(t) indicates air-puff delivery, and *W _airpuff_* is the air-puff penalty. The unchosen-port value was unchanged. Thus, values at each choice reflected outcomes of preceding completed trials. The probability of choosing the right port was calculated with a logistic choice function:

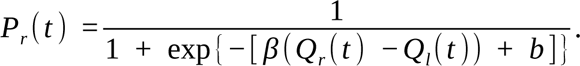

Here, β is the inverse temperature, which determines sensitivity to the right-left value difference, and b is a right-choice bias. The air-puff penalty was fixed at approximately 1.19 relative to a reward value of 1. For each recording, α, β, and b were fitted by minimizing the negative log-likelihood of the observed choices using L-BFGS-B, with α bounded between 0 and 1 and β between 0 and 100. We used six initial parameter settings and retained the converged fit with the lowest negative log-likelihood.

We calculated the right-choice log-odds score, *β* (*Q_r_ −Q_l_*) + *b*, and its logistic transform, P(right). For population analyses, the probability assigned to the currently correct side was P(right) when right was rewarded and 1 − P(right) when left was rewarded.

Calibration compared fitted correct-choice probabilities with observed correct-choice fractions in five equal-width probability bins spanning 0 to 1, calculated within mouse and then averaged with equal weight across mice.

#### M10.6 Continuous signals during return to engagement

To measure transition-related activity without selecting signal peaks, we compared continuous signals during DIS-to-ENG transitions with matched intervals of stable DIS. A transition extended from the retrospectively identified rise onset of the centered 20-s IDF to stable ENG entry. Its duration was mapped to phases 0-1; 10 s of preceding DIS and following ENG were mapped to phases −1-0 and 1-2. Each transition was paired with up to 20 deterministically selected stable-DIS intervals of the same duration from the same session. Session-standardized signals were averaged across DMS and DLS without event-triggered baseline subtraction. The matched stable-DIS mean was then subtracted within transition, session, and mouse. The first initiation after transition onset provided an additional timing anchor. Signal maxima were located after three-bin smoothing within phases 0-1 and averaged within session and mouse.

#### M10.7 Task and movement context of signal events

Each peak above the session’s 97th percentile was assigned the nearest mutually exclusive task-event label within 2 s, or a no-nearby-event label. Movement context was the most frequent B-SOiD label within ±1 s, grouped as task or port behavior, grooming, rearing, and other or unclassified behavior. Expected label fractions were calculated from their prevalence across the full session, weighted by session peak counts.

### M11 Signal features and temporal prediction

These analyses quantified signal representations of behavioral state, progression within a state, and responses around dopamine events. They used the simultaneous recordings in M1.1. Prediction was evaluated on held-out sessions or mice as specified below; imputation, scaling, and learned waveform categories were fitted using training data only.

#### M11.1 Signal features and phase definitions

We summarized fluorescence within each session, region, and state-phase condition using 19 dopamine features, 17 astrocyte features, and 28 features describing their relationship. The 64 measurements and their analytical definitions are listed in Supplementary Table S1. Baseline-normalized signals yielded 1,488 feature vectors (62 sessions × 2 regions × 12 conditions). Each vector summarized windows from the same condition within its session and region.

Two phase divisions served different analyses. For feature summaries, task-action composition, and the response-window placement control, each stable ENG or DIS bout was divided into four quarters labeled C1, C2, C3, and Post, with one-quarter-duration Pre and Post2 flanks outside the bout. For dopamine-aligned response trajectories, sequence analyses, and initiation-aligned comparisons, each stable episode was divided into three equal periods, C1-C3, with exterior Pre and Post flanks of the same duration. Thus, Post lay inside the stable bout for feature summaries and outside it for event trajectories. ENG and DIS trajectories were analyzed separately. An event could appear in two adjacent state-centered trajectories where flanks overlapped, but only once within each trajectory.

#### M11.2 Dimensionality reduction and classification

UMAP (*53*) visualized the organization of each feature family after median imputation and standardization, using Euclidean distance, 20 neighbors, and a minimum distance of 0.25. Statistical predictions used the original feature matrices. Separation of the stable states and displacement of C1 in the relation-feature embedding were measured as distances between centroids divided by within-state spread. Sensitivity analyses varied window duration and embedding initialization.

Stable ENG and DIS were classified from C2/C3 feature vectors using class-balanced logistic regression. Each fold held out one complete recording session. Balanced accuracy and its SEM were calculated across the 62 sessions, and mean accuracy was also calculated within each mouse. The primary statistic was the equal-session mean difference from chance (0.5). A two-sided sign-flip test used 99,999 draws, sharing one sign per session across feature families; the maximum absolute statistic across the three families controlled family-wise error, with plus-one P values.

Early versus core phase classification compared C1 with C2/C3 separately within ENG and DIS, holding out complete mice. Imputation, scaling, and class-balanced logistic regression were fitted within each training fold. Fitting and scoring gave equal total weight to each session and region. The null used 100 label permutations within mouse and region, with P=(1 + number of null accuracies at least as large as observed)/101.

Individual-feature models used the same leave-one-mouse-out approach. Weights were inversely proportional to the number of windows in each mouse-session-region group and were applied in both fitting and scoring. Standalone balanced accuracy measured prediction from one feature with region as a covariate. Unique information was the held-out log-loss improvement of the complete feature family over the family without that feature, expressed in bits per session-region-condition vector. Each vector summarized the corresponding local windows.

#### M11.3 Contributions of current behavior and temporal phase

To quantify the information associated with position in the engagement cycle, ordinary least-squares models predicted dopamine or astrocyte feature vectors in held-out sessions. Predictors were added in sequence: categorical IDF bin, continuous fitted action value, indicators of stable-core state, and state-specific phase categories. Target features were standardized within each training fold. A 4,999-draw control permuted phase order within state and session while preserving IDF, action value, core state, and paired regions.

#### M11.4 Dopamine-aligned astrocyte responses

Transition responses were compared at matched dopamine amplitudes using bins that contained DMS and DLS observations in all three conditions: DIS-to-ENG, stable control, and ENG-to-DIS. The complete paired subset contained 291 of 310 session-bin combinations from 61 sessions and five mice, with 57-59 sessions per amplitude bin. Dopamine magnitudes were averaged equally across regions within session and bin. Each region’s stable astrocyte response was subtracted from its transition response, retaining the regions separately. Linear regressions described the bin means; absolute regional summaries used all available bins. Dopamine increases were measured relative to the preceding 2 s, and transition-minus-stable offsets were evaluated at Δz=1.

Response offsets and slopes were compared within session. A sign-flip test with 99,999 draws shared one sign per mouse-session across regions and metrics. The largest absolute standardized statistic across the eight combinations of transition direction, region, and response metric controlled multiple comparisons. Mouse means summarized session effects.

Changes across the dopamine-aligned astrocyte waveform were sampled in early (1-6 s) and later (10-20 s) windows. These were measurement windows within the evolving response. The later response was the mean signal 10-20 s after the dopamine peak minus the mean during the preceding 2 s. The two windows were analyzed separately and jointly using the equal-third trajectories in M11.1. Phase summaries were averaged within session and then mouse; regions were pooled for trajectory traces and retained in inferential models. Task-action composition controls used the four-quarter definition, whereas phase-dependent gain and initiation-aligned comparisons used thirds and retained DMS and DLS separately.

#### M11.5 Waveform categories and sequence prediction

Waveform categories, termed signatures, grouped dopamine-aligned astrocyte responses with similar preceding activity and early and later response shapes. Eleven waveform features sampled activity before dopamine, at its arrival, and during the early and later response. Effects of nonlinear dopamine amplitude and region were removed before classification. Training sessions defined the categories using a Gaussian mixture with diagonal covariance. Bayesian information criterion selected five or six categories; sensitivity analyses used four to six. A Markov filter represented transitions between categories while allowing variable dwell times.

Sequence models tested whether the current category helped predict the next category beyond exact behavioral phase, position within phase, behavior, dopamine amplitude, and region. Forward sequences were compared with models trained identically on reversed transition laws. Prediction gains were the improvement in log probability assigned to the observed next category, expressed in bits per event; 2 raised to this gain gives the change in geometric-mean assigned probability. Controls used 5,000 circular shifts within session, region, state, and phase.

#### M11.6 Prediction of behavioral boundaries

Boundary models predicted whether the next behavior-defined phase boundary would occur before the next dopamine event. They tested information from the current astrocyte waveform after accounting for behavioral and signal controls available at prediction time, and then the contribution of the preceding waveform sequence beyond the current waveform. Analyses distinguished entry, progression between phases, and exit along ENG and DIS trajectories. Correct temporal alignment was assessed with the within-session, region, state, and phase shifts described in M11.5.

#### M11.7 Prediction from preceding dopamine and astrocyte activity

We tested how much preceding dopamine activity contributed to astrocyte responses beyond current dopamine, movement, and behavioral phase. Dopamine events were detected in baseline-normalized signals; predictors and responses were measured at the same recording site after 180-s rolling-median subtraction. Ridge models predicted early, later, and joint astrocyte responses in held-out sessions. The dataset contained 25,099 assignments of events to state-centered trajectories across 62 sessions, including events shared by adjacent trajectories as defined in M11.1. Models added current local dopamine and movement, categorical state phase, recent dopamine history without state labels, and both phase and history. The history increment after phase measured additional event-to-event information within a phase.

For continuous activity, models predicted the current 1-s astrocyte bin from an accumulated measure of preceding dopamine that ended before that bin. Controls included current dopamine, exact phase, behavior, region, and the astrocyte signal’s own history. Matched multivariate ridge models compared four predictor families: recent dopamine summarized over seven timescales, four current-dopamine features, three measures of astrocyte activity before dopamine, and six measures of the ensuing astrocyte response. Models tested the addition of phase beyond region, followed by phase-specific progress within the phase.

To test whether recent dopamine could reproduce the later ENG astrocyte trajectory, models predicted responses without phase labels. Observed and residual response means across the five ENG phases were centered separately within each mouse. Residual variation was 100 × Σ(residual − mean residual)² / Σ(observed − mean observed)², and the reduction attributed to dopamine history was 100 minus this percentage. Mouse percentages were averaged with SEM.

The contribution of preceding astrocyte activity was tested in separate leave-one-mouse-out models of the 1-6-s response, using signals after 180-s rolling-median subtraction. Baseline predictors included dopamine, behavior, and current and preceding behavioral state. The final step added the astrocyte mean, change, and SD during the preceding 5 s.

### M12 Dopamine-astrocyte coupling

#### M12.1 Event detection and magnitude-dependent responses

To measure how coupling varied with event size, dopamine and astrocyte peaks were detected independently within each session and region after 180-s rolling-median subtraction and session standardization. Astrocyte peaks were grouped by session percentiles into 20-70, 70-90, 90-95, 95-98, 98-99, and 99-100 bands. Dopamine events were baseline-subtracted using −15 to −5 s and ranked by their response at time zero, with bands of 40-70, 70-90, 90-95, 95-98, 98-99, and 99-100. Dopamine magnitude alone determined inclusion from the 40th percentile upward. Response windows measured dopamine during −7 to 0 s and 10 to 30 s around an astrocyte event, and astrocytes during 0 to 7 s after a dopamine event.

Event-conditioned covariance summarized positive lags of +1 to +7 s, when dopamine led astrocytes, and negative lags of −30 to −10 s, when astrocytes led dopamine. High-minus-low contrasts compared the 99th-100th percentile band with the 20th-70th band for astrocyte events or the 40th-70th band for dopamine events. Band means were calculated within session, averaged separately within mouse, and then weighted equally across mice. A null distribution was generated by circularly shifting ordered magnitude labels within session and region while keeping outcomes fixed; permitted shifts had a median temporal displacement of at least 60 s. Each of 100,000 draws used the same averaging procedure. Two-sided centered P values were Holm-adjusted across the 10 outcome-by-region comparisons. Exact Wilcoxon tests across five mouse contrasts and continuous slopes per 10 percentile points provided additional comparisons.

#### M12.2 Coupling after accounting for behavior and signal history

Recent dopamine was represented on a 1-s grid by its signed area relative to the preceding 2-s mean, accumulated with exponential decay over a 10-s time constant. This measure retained both local elevations and suppression. Astrocyte activity was the corresponding 1-s mean after 180-s rolling-median subtraction and session standardization. Bins were included when both the anchor and prediction target fell within the behavior-defined ENG or DIS trajectories, including Pre and Post flanks.

To estimate coupling beyond shared behavior and each signal’s own history, we separately removed these contributions from dopamine history and astrocyte activity within session and region. Predictors were state and phase, progress within phase, IDF and its derivative, movement speed, initiation rate, and each signal’s preceding values at 1, 2, 5, 10, and 20 s. Cross-correlations were then calculated from the residuals, with positive lag indicating dopamine leading astrocytes. A 1,000-permutation control shifted astrocyte residuals independently within session and region by at least 60 s. The dopamine-leading component was the maximum from +1 to +10 s; the astrocyte-leading component was the mean from −20 to −5 s. Regional curves were averaged across sessions within each mouse before measuring these components.

#### M12.3 Prediction in both signal directions

Matched models tested whether each signal’s history improved prediction of the other signal in held-out sessions. Both directions used the same 1-s bins, prediction horizons of 1, 2, 5, 10, 20, and 40 s, and history lags of 0, 1, 2, 5, 10, and 20 s. The base model included region, the target signal’s own history, and current and target-time state, phase, progress within phase, IDF and its derivative, movement speed, and initiation rate. The comparison model added the other signal’s history. Improvement was the reduction in held-session mean-squared error, averaged within mouse and then across five mice. Standardized coefficients for the added history terms were summed within each outer fold and averaged across folds to determine the sign of the cross-signal relationship.

#### M12.4 Event-selection and recording-level controls

Detection controls selected dopamine and astrocyte peaks independently above their session-specific 97th percentiles, with minimum separations of 1 and 2 s, respectively. Nearest opposite-signal timing was calculated without changing either anchor set. Event-aligned responses used the fixed windows in M12.1. A separate analysis retained only the strongest event in each 6-s cluster. Session estimates were averaged within mouse, and exact two-sided Wilcoxon tests were Holm-adjusted across the prespecified event-response comparisons.

For recording-level prevalence, whole-session cross-correlations were calculated separately for DMS and DLS. We selected the dopamine-leading maximum from +1 to +7 s and the astrocyte-leading minimum from −30 to −10 s, then calculated the fraction of recordings containing both expected signs.

### M13 Analysis of optogenetic responses

#### M13.1 Response magnitudes and timing

Responses to SNpc dopamine-neuron stimulation were analyzed in separate dopamine, astrocyte-calcium, and adenosine recording cohorts using the protocol in M2.1. Astrocytic receptor-deletion cohorts and the adenosine PMCA cohort were compared with their matched controls. Response magnitudes and time courses were summarized within mouse and across mice as means with SEM. Receptor-deletion and adenosine-PMCA comparisons used two-sided Mann-Whitney U tests.

#### M13.2 Adaptation and recovery between stimulation trains

Responses across ten repeated trains were summarized within mouse before comparison across animals. Linear and exponential declines were fitted separately for each mouse, region, and stimulation frequency. We calculated ΔAIC as AIClinear − AICexponential; values above 2 favored an exponential decline. There were 14 astrocyte and 16 dopamine fits. In paired-train experiments, the second response was divided by the first for each inter-train interval and region, and regional means and SEM were calculated from mouse values.

#### M13.3 Responses to astrocyte stimulation

Direct astrocyte stimulation used the protocol in M2.2, with four calcium-recording and five dopamine-recording mice. Phase-averaged responses were tested against zero with one-sample t tests; timing was compared between cohorts with two-sided Mann-Whitney U tests. Dopamine recovery was measured as the time when suppression declined to within 15% of its peak magnitude, the first re-entry into the baseline band, and sustained return to that band. These endpoints measured partial recovery, first crossing, and sustained return, respectively. The open-field assay and movement analyses are described in M6.2 and M7.

### M14 Discrimination, engagement, and operant performance

#### M14.1 Discrimination and response propensity

Reward- and cost-cue response rates were calculated from licking in the designated pre-outcome response window of the approach-avoidance task (M6.3). Discrimination was d′ = z(hit rate) − z(false-alarm rate), and response criterion was C = −[z(hit rate) + z(false-alarm rate)]/2. Engagement was represented by −C. We compared engagement at matched discrimination and discrimination at matched engagement.

#### M14.2 Comparisons across learning stages

The action-selectivity trajectory was the arithmetic mean of the control and PMCA group means at each d′ position. It located the behavioral change, while the prespecified d′ interval of 0.5-0.75 defined the primary transition-stage comparison. All univariate group P values used exhaustive two-sided permutation of mouse labels. The two cue-response comparisons at the transition were Holm-adjusted together.

To test whether the engagement deficit was concentrated at the transition, we calculated transition engagement minus the average of low- and established-stage engagement for each mouse with all three stages, then permuted group labels. An overlapping-window analysis used the maximum cluster mass across the full trajectory under mouse-label permutation.

#### M14.3 Joint cue responses and licking controls

Joint reward-cost response comparisons measured the Mahalanobis distance between group centroids. Exact P values were the fraction of all mouse-label allocations, preserving group sizes, with a distance at least as large as observed. Comparisons were made at low, transition, and established discrimination and at low, middle, and high engagement.

Response-lick distributions and consummatory licks per reward were summarized within mouse and compared by mouse-label permutation. The fraction of mice exceeding the prespecified maximum-session d′ threshold of 1.2 was compared with a two-sided Fisher exact test.

#### M14.4 Prior operant performance

The prior-task analysis used the same eight GFAP-Cre-injected control and seven PMCA mice later tested in the approach-avoidance task. Six prespecified endpoints summarized late fixed-ratio-1 reward rate, progressive-ratio breakpoint, no-lick-gated task cycles and completion time, and late variable-interval hit rate and cue selectivity. Group comparisons used exact two-sided mouse-label permutation. Confidence intervals were obtained by resampling mice within group. Benjamini-Hochberg correction was applied jointly across all 18 prior-task group-comparison endpoints, including the six primary performance endpoints and fixed-ratio-1 licking microstructure. Cue-directed licking was measured over the first 2 s after cue onset and divided by the 2-s window duration to obtain Hz. Intertrial licking was also expressed in Hz. Cue selectivity was calculated from these rates; hit rates were calculated as the fraction of completed cue trials with a hit.

### M15 Software and statistical reporting

Analyses were performed in Python using SciPy 1.15.2 and statsmodels 0.14.6 for signal processing and statistics, scikit-learn 1.2.2 for classification and regression, and umap-learn 0.5.12 for UMAP. InceptionTime used tsai 0.4.1 with PyTorch 2.7.1 and fastai 2.8.7; MiniROCKET used tsai 0.4.0. Complete software dependencies and implementation details accompany the released analysis code.

Mouse was the independent biological unit for comparisons between experimental groups. Repeated sessions, trials, bouts, and signal events were summarized as specified for each analysis. Unless a directional test was prespecified, tests were two-sided. Randomization units, weighting, and multiple-comparison corrections are given with the relevant procedures; sample sizes and displayed summaries are reported in the figure legends or statistics table. References are listed in the main manuscript bibliography.

#### AI use

GPT-6 Astra (OpenAI; model identifier: gpt-6-astra), accessed through ChatGPT and Codex, was used to assist with reviewing and debugging analysis code. The authors take full responsibility for the final code and analyses.

**Table M1.** Viral constructs and source records. Table M1 lists the constructs and catalog identifiers, with originating publications or construct records for vectors, sensors, and actuators. Shared constructs are listed once; Drd1 and Drd2 identifiers refer to the plasmids.

| Construct and use | Serotype | Supplier / catalog | Publication / construct record |
| --- | --- | --- | --- |
| Syn-FLEX-rc[ChrimsonR-tdTomato] Dopamine-neuron stimulation (M2.1) | AAV5 | Addgene #62723 | Klapoetke et al., Nature Methods, 2014 (39). Exact deposited construct. |
| hSyn-DA3h-WPRE-hGH (green DA3h) Dopamine recording (M2.1-M2.2) | Not reported | BrainVTA #PT-4721 | Zhuo et al., Nature Methods, 2024 (40). Sensor-development paper. |
| gfaABC1D-cyto-GCaMP6f Astrocyte calcium recording (M2.1-M2.3) | AAV5 | Addgene #52925 | Haustein et al., Neuron, 2014 (54): exact astrocyte vector. Chen et al., Nature, 2013 (41): GCaMP6 indicator. |
| GFAP-Cre / pAAV.GFAP.Cre.WPRE.hGH Astrocyte targeting, AP/AV control, and receptor deletion (M2.2, M2.4-M2.5) | AAV5 | Addgene #105550-AAV5 | Unpublished deposit; James M. Wilson laboratory (Addgene record). |
| nEF-Con/Foff 2.0-ChRmine-oScarlet Astrocyte stimulation (M2.2) | AAV8 | Addgene #137161-AAV8 | Fenno et al., Neuron, 2020 (43): exact intersectional vector. Marshel et al., Science, 2019 (42): ChRmine actuator. |
| hSyn-GRAB-rDA3h (red rDA3h) Simultaneous dopamine/calcium recording (M2.3) | AAV2/9 | BrainVTA #PT-5198 | Zhuo et al., Nature Methods, 2024 (40). Sensor-development paper. |
| GfaABC1D-mCherry-hPMCA2w/b (CalEx) Astrocyte calcium attenuation (M2.4; M13) | AAV5 | Addgene #111568-AAV5 | Yu et al., Neuron, 2018 (21). CalEx construct. |
| hSyn-Ado_B10 (Ado1.0)-WPRE-hGH polyA Adenosine recording (M2.5; M13) | AAV2/8 | BrainVTA #PT-1348 | BrainVTA Ado_B10 (Ado1.0) construct record. |
| CMV-FLEX-SaCas9-U6-sgDrd1 Astrocytic Drd1 deletion (M2.5) | AAV5 | Addgene plasmid #159917 | Unpublished deposit; Larry Zweifel laboratory (Addgene record). |
| CMV-FLEX-SaCas9-U6-sgDrd2 Astrocytic Drd2 deletion (M2.5) | AAV5 | Addgene plasmid #159918 | Unpublished deposit; Larry Zweifel laboratory (Addgene record). |

Catalog records: Addgene #62723; Addgene #52925; Addgene #105550; Addgene #137161; Addgene #111568; Addgene plasmid #159917; Addgene plasmid #159918; BrainVTA #PT-1348

## Acknowledgments

We thank Henry Hall for laboratory management and Dr. Yasuo Kubota for submission management.

## Author contributions

- IL and AMG conceptualized and designed the study, interpreted the results, and wrote the manuscript.
- IL designed the experimental procedures, performed experiments, and analyzed the data.
- EH designed and conducted the approach-avoidance experiments and analyzed their results.
- GA and WC contributed to the development of preprocessing code and data analyses.
- KH developed the kinematic analyses.
- SD developed the convolutional neural network analyses.

## Data, code, and materials availability

The data and analysis code supporting this study will be publicly available through Zenodo upon journal publication. No new biological materials were generated. Users are requested to cite the associated paper when reusing these resources.

## AI use

GPT-6 Astra (OpenAI; model identifier: gpt-6-astra), accessed through ChatGPT and Codex, was used during manuscript preparation to assist with proofreading and refining manuscript text for clarity and conciseness and with reviewing and debugging code. The authors take full responsibility for the accuracy and integrity of the final manuscript.

## References and Notes

1. D. J. Surmeier, J. Ding, M. Day, Z. Wang, W. Shen, D1 and D2 dopamine-receptor modulation of striatal glutamatergic signaling in striatal medium spiny neurons. Trends in Neurosciences 30, 228–235 (2007). 10.1016/j.tins.2007.03.008.

2. J. Cox, I. B. Witten, Striatal circuits for reward learning and decision-making. Nature Reviews Neuroscience 20, 482–494 (2019). 10.1038/s41583-019-0189-2.

3. A. M. Graybiel, The Basal Ganglia and Chunking of Action Repertoires. Neurobiology of Learning and Memory 70, 119–136 (1998). 10.1006/nlme.1998.3843.

4. A. M. Graybiel, A. Matsushima, Striosomes and Matrisomes: Scaffolds for Dynamic Coupling of Volition and Action. Annual Review of Neuroscience 46, 359–380 (2023). 10.1146/annurev-neuro-121522-025740.

5. M. W. Howe, D. A. Dombeck, Rapid signalling in distinct dopaminergic axons during locomotion and reward. Nature 535, 505–510 (2016). 10.1038/nature18942.

6. A. Mohebi, J. R. Pettibone, A. A. Hamid, J.-M. T. Wong, L. T. Vinson, T. Patriarchi, L. Tian, R. T. Kennedy, J. D. Berke, Dissociable dopamine dynamics for learning and motivation. Nature 570, 65–70 (2019). 10.1038/s41586-019-1235-y.

7. M. Corkrum, A. Covelo, J. Lines, L. Bellocchio, M. Pisansky, K. Loke, R. Quintana, P. E. Rothwell, R. Lujan, G. Marsicano, E. D. Martin, M. J. Thomas, P. Kofuji, A. Araque, Dopamine-Evoked Synaptic Regulation in the Nucleus Accumbens Requires Astrocyte Activity. Neuron 105, 1036–1047.e5 (2020). 10.1016/j.neuron.2019.12.026.

8. M. Corkrum, A. Araque, Astrocyte-neuron signaling in the mesolimbic dopamine system: the hidden stars of dopamine signaling. Neuropsychopharmacology 46, 1864–1872 (2021). 10.1038/s41386-021-01090-7.

9. B. S. Khakh, Astrocyte-Neuron Interactions in the Striatum: Insights on Identity, Form, and Function. Trends in Neurosciences 42, 617–630 (2019). 10.1016/j.tins.2019.06.003.

10. F. Ding, J. O’Donnell, A. S. Thrane, D. Zeppenfeld, H. Kang, L. Xie, F. Wang, M. Nedergaard, α1-Adrenergic receptors mediate coordinated Ca2+ signaling of cortical astrocytes in awake, behaving mice. Cell Calcium 54, 387–394 (2013). 10.1016/j.ceca.2013.09.001.

11. K. B. Lefton, Y. Wu, Y. Dai, T. Okuda, Y. Zhang, A. Yen, G. M. Rurak, S. Walsh, R. Manno, B. E. Myagmar, J. D. Dougherty, V. K. Samineni, P. C. Simpson, T. Papouin, Norepinephrine signals through astrocytes to modulate synapses. Science 388, 776–783 (2025). 10.1126/science.adq5480.

12. A. B. Chen, M. Duque, A. Rymbek, M. Dhanasekar, V. M. Wang, X. Mi, L. Tocquer, S. Narayan, E. Marquez Legorreta, M. Eddison, G. Yu, C. Wyart, D. A. Prober, F. Engert, M. B. Ahrens, Norepinephrine changes behavioral state through astroglial purinergic signaling. Science 388, 769–775 (2025). 10.1126/science.adq5233.

13. K. E. Poskanzer, R. Yuste, Astrocytes regulate cortical state switching in vivo. Proceedings of the National Academy of Sciences 113, E2675–E2684 (2016). 10.1073/pnas.1520759113.

14. E. Montalban, A. Ansoult, D. Herrera Moro Chao, C. Pham, C. Franco, A. Contini, J. Castel, R. Hassouna, M. H. Hardonk, A. Petitbon, E. Foppen, G. Gangarossa, P. Trifilieff, D. Li, S. Luquet, C. Martin, Striatal astrocytes modulate behavioral flexibility and whole-body metabolism in mice. Nature Communications 16, 5417 (2025). 10.1038/s41467-025-60968-y.

15. L.-H. Tai, A. M. Lee, N. Benavidez, A. Bonci, L. Wilbrecht, Transient stimulation of distinct subpopulations of striatal neurons mimics changes in action value. Nature Neuroscience 15, 1281–1289 (2012). 10.1038/nn.3188.

16. I. Lazaridis, J. R. Crittenden, G. Ahn, K. Hirokane, I. R. Wickersham, T. Yoshida, A. Mahar, V. Skara, J. H. Loftus, K. Parvataneni, K. Meletis, J. T. Ting, E. Hueske, A. Matsushima, A. M. Graybiel, Striosomes control dopamine via dual pathways paralleling canonical basal ganglia circuits. Current Biology 34, 5263–5283.e8 (2024). 10.1016/j.cub.2024.09.070.

17. B. M. Roberts, E. Lambert, J. A. Livesey, Z. Wu, Y. Li, S. J. Cragg, Dopamine Release in Nucleus Accumbens Is under Tonic Inhibition by Adenosine A1 Receptors Regulated by Astrocytic ENT1 and Dysregulated by Ethanol. Journal of Neuroscience 42, 1738–1751 (2022). 10.1523/JNEUROSCI.1548-21.2021.

18. S. Kang, S.-I. Hong, J. Lee, L. Peyton, M. Baker, S. Choi, H. Kim, S.-Y. Chang, D.-S. Choi, Activation of Astrocytes in the Dorsomedial Striatum Facilitates Transition From Habitual to Goal-Directed Reward-Seeking Behavior. Biological Psychiatry 88, 797–808 (2020). 10.1016/j.biopsych.2020.04.023.

19. M. L. Cooper, M. C. Selles, M. Cammer, C. Redd, H. K. Gildea, J. Sall, K. E. Chiurri, P. Cheung, D. G. Wheeler, A. S. Saab, S. A. Liddelow, M. V. Chao, Astrocytes connect specific brain regions through plastic networks. Nature 655, 183–191 (2026). 10.1038/s41586-026-10426-6.

20. L. Bojarskaite, D. M. Bjørnstad, K. H. Pettersen, C. Cunen, G. H. Hermansen, K. S. Åbjørsbråten, A. R. Chambers, R. Sprengel, K. Vervaeke, W. Tang, R. Enger, E. A. Nagelhus, Astrocytic Ca2+ signaling is reduced during sleep and is involved in the regulation of slow wave sleep. Nature Communications 11, 3240 (2020). 10.1038/s41467-020-17062-2.

21. X. Yu, A. M. W. Taylor, J. Nagai, P. Golshani, C. J. Evans, G. Coppola, B. S. Khakh, Reducing Astrocyte Calcium Signaling In Vivo Alters Striatal Microcircuits and Causes Repetitive Behavior. Neuron 99, 1170–1187.e9 (2018). 10.1016/j.neuron.2018.08.015.

22. A. Berditchevskaia, R. D. Cazé, S. R. Schultz, Performance in a GO/NOGO perceptual task reflects a balance between impulsive and instrumental components of behaviour. Scientific Reports 6, 27389 (2016). 10.1038/srep27389.

23. K. A. Guttenplan, I. Maxwell, E. Santos, L. A. Borchardt, E. Manzo, L. Abalde-Atristain, R. D. Kim, M. R. Freeman, GPCR signaling gates astrocyte responsiveness to neurotransmitters and control of neuronal activity. Science 388, 763–768 (2025). 10.1126/science.adq5729.

24. S. N. Schiffmann, J. J. Vanderhaeghen, Adenosine A2 receptors regulate the gene expression of striatopallidal and striatonigral neurons. Journal of Neuroscience 13, 1080–1087 (1993). 10.1523/JNEUROSCI.13-03-01080.1993.

25. C. Cervetto, G. Maura, D. Guidolin, S. Amato, C. Ceccoli, L. F. Agnati, M. Marcoli, Striatal astrocytic A2A-D2 receptor-receptor interactions and their role in neuropsychiatric disorders. Neuropharmacology 237, 109636 (2023). 10.1016/j.neuropharm.2023.109636.

26. S. W. Schoen, A. M. Graybiel, Species-specific patterns of glycoprotein expression in the developing rodent caudoputamen: association of 5′-nucleotidase activity with dopamine islands and striosomes in rat, but with extrastriosomal matrix in mouse. Journal of Comparative Neurology 333, 578–596 (1993). 10.1002/cne.903330410.

27. P. Albulescu, I. Macsinga, A. Rusu, C. Sulea, A. Bodnaru, B. T. Tulbure, "Give me a break!" A systematic review and meta-analysis on the efficacy of micro-breaks for increasing well-being and performance. PLOS ONE 17, e0272460 (2022). 10.1371/journal.pone.0272460.

28. A. Ariga, A. Lleras, Brief and rare mental "breaks" keep you focused: Deactivation and reactivation of task goals preempt vigilance decrements. Cognition 118, 439–443 (2011). 10.1016/j.cognition.2010.12.007.

29. M. Bönstrup, I. Iturrate, R. Thompson, G. Cruciani, N. Censor, L. G. Cohen, A Rapid Form of Offline Consolidation in Skill Learning. Current Biology 29, 1346–1351.e4 (2019). 10.1016/j.cub.2019.02.049.

30. E. R. Buch, L. Claudino, R. Quentin, M. Bönstrup, L. G. Cohen, Consolidation of human skill linked to waking hippocampo-neocortical replay. Cell Reports 35, 109193 (2021). 10.1016/j.celrep.2021.109193.

31. X. Jin, R. M. Costa, Start/stop signals emerge in nigrostriatal circuits during sequence learning. Nature 466, 457–462 (2010). 10.1038/nature09263.

32. M. S. Jog, Y. Kubota, C. I. Connolly, V. Hillegaart, A. M. Graybiel, Building Neural Representations of Habits. Science 286, 1745–1749 (1999). 10.1126/science.286.5445.1745.

33. K. S. Smith, A. M. Graybiel, A Dual Operator View of Habitual Behavior Reflecting Cortical and Striatal Dynamics. Neuron 79, 361–374 (2013). 10.1016/j.neuron.2013.05.038.

34. M. Brancaccio, A. P. Patton, J. E. Chesham, E. S. Maywood, M. H. Hastings, Astrocytes Control Circadian Timekeeping in the Suprachiasmatic Nucleus via Glutamatergic Signaling. Neuron 93, 1420–1435.e5 (2017). 10.1016/j.neuron.2017.02.030.

35. A. González-Vila, M. Luengo-Mateos, M. Silveira-Loureiro, P. Garrido-Gil, N. Ohinska, M. González-Domínguez, J. L. Labandeira-García, C. García-Cáceres, M. López, O. Barca-Mayo, Astrocytic insulin receptor controls circadian behavior via dopamine signaling in a sexually dimorphic manner. Nature Communications 14, 8175 (2023). 10.1038/s41467-023-44039-8.

36. V. Untiet, F. R. M. Beinlich, P. Kusk, N. Kang, A. Ladrón-de-Guevara, W. Song, C. Kjaerby, M. Andersen, N. Hauglund, Z. Bojarowska, B. Sigurdsson, S. Deng, H. Hirase, N. C. Petersen, A. Verkhratsky, M. Nedergaard, Astrocytic chloride is brain state dependent and modulates inhibitory neurotransmission in mice. Nature Communications 14, 1871 (2023). 10.1038/s41467-023-37433-9.

37. S. M. Theparambil, O. Kopach, A. Braga, S. Nizari, P. S. Hosford, V. Sagi-Kiss, A. Hadjihambi, C. Konstantinou, N. Esteras, A. Gutierrez Del Arroyo, G. L. Ackland, A. G. Teschemacher, N. Dale, T. Eckle, P. Andrikopoulos, D. A. Rusakov, S. Kasparov, A. V. Gourine, Adenosine signalling to astrocytes coordinates brain metabolism and function. Nature 632, 139–146 (2024). 10.1038/s41586-024-07611-w.

38. A. M. Graybiel, Habits, Rituals, and the Evaluative Brain. Annual Review of Neuroscience 31, 359–387 (2008). 10.1146/annurev.neuro.29.051605.112851.

## References cited only in Materials and Methods

39. N. C. Klapoetke, Y. Murata, S. S. Kim, S. R. Pulver, A. Birdsey-Benson, Y. K. Cho, T. K. Morimoto, A. S. Chuong, E. J. Carpenter, Z. Tian, J. Wang, Y. Xie, Z. Yan, Y. Zhang, B. Y. Chow, B. Surek, M. Melkonian, V. Jayaraman, M. Constantine-Paton, G. K.-S. Wong, E. S. Boyden, Independent optical excitation of distinct neural populations. Nature Methods 11, 338–346 (2014). 10.1038/nmeth.2836.

40. Y. Zhuo, B. Luo, X. Yi, H. Dong, X. Miao, J. Wan, J. T. Williams, M. G. Campbell, R. Cai, T. Qian, F. Li, S. J. Weber, L. Wang, B. Li, Y. Wei, G. Li, H. Wang, Y. Zheng, Y. Zhao, M. E. Wolf, Y. Zhu, M. Watabe-Uchida, Y. Li, Improved green and red GRAB sensors for monitoring dopaminergic activity in vivo. Nature Methods 21, 680–691 (2024). 10.1038/s41592-023-02100-w.

41. T. W. Chen, T. J. Wardill, Y. Sun, S. R. Pulver, S. L. Renninger, A. Baohan, E. R. Schreiter, R. A. Kerr, M. B. Orger, V. Jayaraman, L. L. Looger, K. Svoboda, D. S. Kim, Ultrasensitive fluorescent proteins for imaging neuronal activity. Nature 499, 295–300 (2013). 10.1038/nature12354.

42. J. H. Marshel, Y. S. Kim, T. A. Machado, S. Quirin, B. Benson, J. Kadmon, C. Raja, A. Chibukhchyan, C. Ramakrishnan, M. Inoue, J. C. Shane, D. J. McKnight, S. Yoshizawa, H. E. Kato, S. Ganguli, K. Deisseroth, Cortical layer-specific critical dynamics triggering perception. Science 365, eaaw5202 (2019). 10.1126/science.aaw5202.

43. L. E. Fenno, C. Ramakrishnan, Y. S. Kim, K. E. Evans, M. Lo, S. Vesuna, M. Inoue, K. Y. M. Cheung, E. Yuen, N. Pichamoorthy, A. S. O. Hong, K. Deisseroth, Comprehensive Dual- and Triple-Feature Intersectional Single-Vector Delivery of Diverse Functional Payloads to Cells of Behaving Mammals. Neuron 107, 836–853.e11 (2020). 10.1016/j.neuron.2020.06.003.

44. A. Mathis, P. Mamidanna, K. M. Cury, T. Abe, V. N. Murthy, M. W. Mathis, M. Bethge, DeepLabCut: markerless pose estimation of user-defined body parts with deep learning. Nature Neuroscience 21, 1281–1289 (2018). 10.1038/s41593-018-0209-y.

45. G. Lopes, N. Bonacchi, J. Frazão, J. P. Neto, B. V. Atallah, S. Soares, L. Moreira, S. Matias, P. M. Itskov, P. A. Correia, R. E. Medina, L. Calcaterra, E. Dreosti, J. J. Paton, A. R. Kampff, Bonsai: an event-based framework for processing and controlling data streams. Frontiers in Neuroinformatics 9, 7 (2015). 10.3389/fninf.2015.00007.

46. T. N. Lerner, C. Shilyansky, T. J. Davidson, K. E. Evans, K. T. Beier, K. A. Zalocusky, A. K. Crow, R. C. Malenka, L. Luo, R. Tomer, K. Deisseroth, Intact-Brain Analyses Reveal Distinct Information Carried by SNc Dopamine Subcircuits. Cell 162, 635–647 (2015). 10.1016/j.cell.2015.07.014.

47. A. I. Hsu, E. A. Yttri, B-SOiD, an open-source unsupervised algorithm for identification and fast prediction of behaviors. Nature Communications 12, 5188 (2021). 10.1038/s41467-021-25420-x.

48. J. F. Tillmann, A. I. Hsu, M. K. Schwarz, E. A. Yttri, A-SOiD, an active-learning platform for expert-guided, data-efficient discovery of behavior. Nature Methods 21, 703–711 (2024). 10.1038/s41592-024-02200-1.

49. H. Ismail Fawaz, B. Lucas, G. Forestier, C. Pelletier, D. F. Schmidt, J. Weber, G. I. Webb, L. Idoumghar, P.- A. Muller, F. Petitjean, InceptionTime: Finding AlexNet for time series classification. Data Mining and Knowledge Discovery 34, 1936–1962 (2020). 10.1007/s10618-020-00710-y.

50. I. Oguiza, tsai [software], version 0.4.1 for CNN and 0.4.0 for MiniROCKET, GitHub (2023); https://github.com/timeseriesAI/tsai (accessed 6 October 2026).

51. A. Dempster, D. F. Schmidt, G. I. Webb, “MiniRocket: A Very Fast (Almost) Deterministic Transform for Time Series Classification” in Proceedings of the 27th ACM SIGKDD Conference on Knowledge Discovery & Data Mining (ACM, 2021), pp. 248-257. 10.1145/3447548.3467231.

52. R. S. Sutton, A. G. Barto, Reinforcement Learning: An Introduction (MIT Press, ed. 2, 2018). https://mitpress.mit.edu/9780262039246/reinforcement-learning/.

53. L. McInnes, J. Healy, N. Saul, L. Großberger, UMAP: Uniform Manifold Approximation and Projection. Journal of Open Source Software 3, 861 (2018). 10.21105/joss.00861.

54. M. D. Haustein, S. Kracun, X.-H. Lu, T. Shih, O. Jackson-Weaver, X. Tong, J. Xu, X. W. Yang, T. J. O’Dell, J. S. Marvin, M. H. Ellisman, E. A. Bushong, L. L. Looger, B. S. Khakh, Conditions and Constraints for Astrocyte Calcium Signaling in the Hippocampal Mossy Fiber Pathway. Neuron 82, 413–429 (2014). 10.1016/j.neuron.2014.02.041.

